# Ataxin-2 polyglutamine expansions aberrantly sequester TDP-43, drive ribonucleoprotein condensate transport dysfunction and suppress local translation

**DOI:** 10.1101/2023.01.30.526372

**Authors:** Denethi Wijegunawardana, Sonali S. Vishal, Neha Venkatesh, Pallavi P. Gopal

## Abstract

Altered RNA metabolism is a common pathogenic mechanism linked to familial and sporadic Amyotrophic lateral sclerosis (ALS). ALS is characterized by mislocalization and aggregation of TDP-43, an RNA-binding protein (RBP) with multiple roles in post-transcriptional RNA processing. Recent studies have identified genetic interactions between TDP-43 and Ataxin-2, a polyglutamine (polyQ) RBP in which intermediate length polyQ expansions confer increased ALS risk. Here, we used live-cell confocal imaging, photobleaching and translation reporter assays to study the localization, transport dynamics and mRNA regulatory functions of TDP-43/Ataxin-2 in rodent primary cortical neurons. We show that Ataxin-2 polyQ expansions aberrantly sequester TDP-43 within ribonucleoprotein (RNP) condensates, and disrupt both its motility along the axon and liquid-like properties. Our data suggest that Ataxin-2 governs motility and translation of neuronal RNP condensates and that Ataxin-2 polyQ expansions fundamentally perturb spatial localization of mRNA and suppress local translation. Overall, these results indicate Ataxin-2 polyQ expansions have detrimental effects on stability, localization, and translation of transcripts critical for axonal and cytoskeletal integrity, particularly important for motor neurons.

## INTRODUCTION

Amyotrophic lateral sclerosis (ALS) and frontotemporal dementia (FTD) are profoundly debilitating and fatal neurodegenerative diseases with overlapping clinical, pathologic and genetic features (1). Despite advances in our understanding of the genetic basis of ALS/FTD, the cellular mechanisms underlying neurodegeneration remain poorly understood. Current treatments are unable to significantly alter disease progression and extend life for only a few months (2)

Genetic, pathologic, and molecular studies strongly implicate TDP-43 (transactive response <u>D</u>NA-binding <u>p</u>rotein of <u>43</u> kDa), a highly conserved, ubiquitously expressed RNA and DNA-binding protein in the pathogenesis of ALS/FTD (3, 4). TDP-43 is essential for pre-mRNA splicing, stabilization and localization of mRNA transcripts, many of which encode synaptic and axonal proteins (5-8). The vast majority (>95%) of ALS patients and nearly half of FTD patients have pathologic aggregates composed of TDP-43 and concomitant loss of TDP-43 from the nucleus, indicating that mis-regulation of TDP-43 is a defining feature of these diseases (3). Furthermore, highly penetrant dominant mutations of the gene encoding TDP-43, *TARDBP*, and other RNA binding proteins highlight altered RNA metabolism as a common pathogenic mechanism in ALS/FTD (9-14).

Multiple studies, from yeast to mammals, demonstrate that TDP-43 interacts genetically with Ataxin-2, a highly conserved polyglutamine (polyQ) and RNA-binding protein previously linked to Spinocerebellar ataxia type 2 (SCA2) (15-20). Ataxin-2 is a member of the Like-Sm (LSm) protein family and is comprised of an N-terminal polyQ tract, globular LSm and LSm-associated domain (LSmAD) RNA-binding domains, as well as a C-terminal <u>p</u>oly(<u>A</u>)-binding protein-interacting motif (PAM2) (21, 22); Fig. 1A. Ataxin-2 enhances TDP-43-induced neurodegeneration in an RNA-dependent manner in a *Drosophila* model of motor neuron disease (15, 16). Conversely, reducing Ataxin-2 levels improves motor function and survival in a TDP-43 transgenic mouse model (17). Moreover, human genetics data and meta-analyses of several ALS patient and control populations worldwide show that the Ataxin-2 polyQ tract, which is 22-23 glutamines (range 16-27) in length in healthy individuals, is more often expanded (29-33 glutamines) in ALS patients (23); Fig. 1A. Additional independent, smaller studies of patients with familial ALS/FTD suggest that the frequency of Ataxin-2 intermediate polyQ repeat expansions also is increased in FTD patients with symptoms of motor neuron disease (FTD-MND), but not in patients with a pure FTD phenotype (24, 25). Together, these data strongly suggest that intermediate-length polyQ expansions in human Ataxin-2 (ATXN2) gene significantly increase risk for ALS and possibly also FTD-MND (16, 24). These data have spurred efforts to reduce Ataxin-2 levels as a new therapeutic strategy in ALS. However, an appreciation of the physiologic role of TDP-43 / Ataxin-2 interactions in neurons and a mechanistic understanding of how Ataxin-2 polyQ expansions increase risk of ALS, are lacking. This knowledge would stimulate development of additional molecular therapies and may also reveal why motor neurons, but not cortical neurons, show particular sensitivity to Ataxin-2 polyQ expansions.

**Figure 1.**
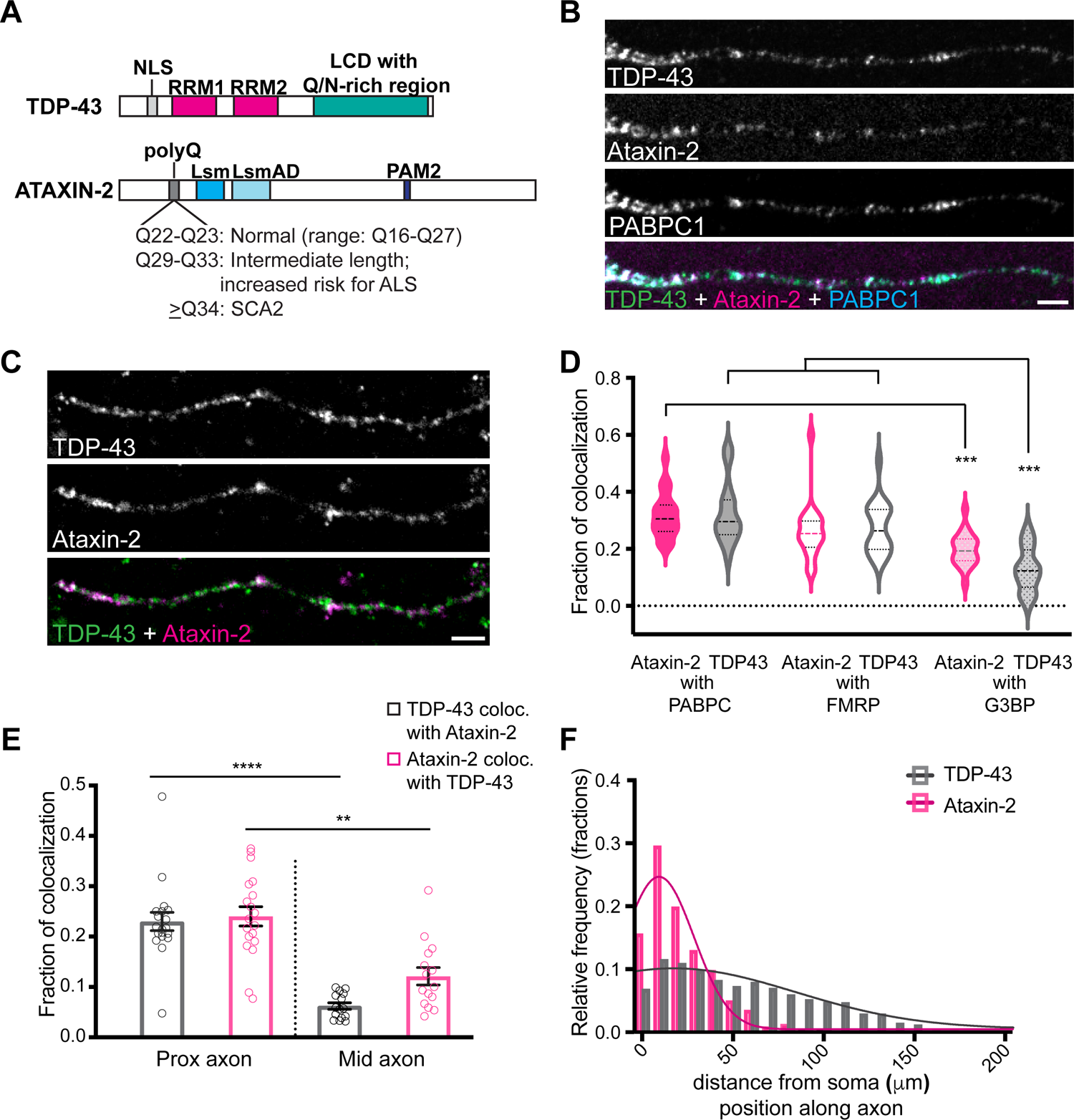
Axonal distribution and colocalization of endogenous TDP-43 and Ataxin-2 in RNP condensates. **(A)** TDP-43 is comprised of an N-terminal domain which contains a nuclear localization signal (NLS); two RNA recognition motif domains (RRM1, RRM2); and a predominantly disordered C-terminal low complexity domain (LCD). Ataxin-2 contains an N-terminal polyglutamine tract that is normally 22-23 glutamines in length; globular RNA-binding domains, Lsm and LsmAD, and a poly(A)-binding protein interacting motif (PAM2). Intermediate-length polyglutamine expansions between Q29-33 are associated with increased risk for ALS, whereas expansions greater than Q34 cause SCA2. **(B, C)** Immunofluorescence for endogenous TDP-43 and Ataxin-2 reveals colocalization in a subset of RNP granules and with neuronal RNP markers (e.g. PABPC1) in mouse cortical neurons (DIV 10-12); Scale bar = 3μm. **(D)** Fraction of TDP-43 or Ataxin-2 that colocalize with each RNP condensate marker (PABPC, FMRP, or G3BP1) in the axon. Quantification by a stringent object-based colocalization method shows that TDP-43 and Ataxin-2 exhibit a higher degree of colocalization with PABPC and FMRP than stress granule marker, G3BP1. (***p *≤* 0.0003, Kruskal-Wallis; *n*=21 neurons per condition, *N* = 3). **(E)** TDP-43 and Ataxin-2 colocalize with each other to a greater extent in the proximal axon than in the mid axon (****p<0.0001 and **p < 0.002, Kruskal-Wallis; *n*=19 neurons for proximal axon, *n*=15 neurons for mid axon, *N* = 3). **(F)** Histogram of the position of Ataxin-2 or TDP-43 condensates along the axon, binned by distance (*μ*m) from the soma. Ataxin-2 is enriched proximally while TDP-43 is distributed throughout the axon (p<0.0001, Kolmogorov-Smirnov; n=536 Ataxin-2 granules, n=1556 TDP-43 granules, from at least n=15 axons each, *N* = 3).

Several lines of evidence suggest physiologic roles for TDP-43 and Ataxin-2 in regulating stability and/or translation of target mRNAs. Ataxin-2 associates with polyribosomes, and PAR-CLIP data reveal direct binding of Ataxin-2 LSm domain to AU- and uridine-rich elements within the 3′ UTRs of target RNAs. Furthermore, Ataxin-2 enhances stability and translation of target mRNAs in a poly(A)-binding protein 1-independent manner (26, 27). In neurons, *Drosophila* Ataxin-2 (dAtx) is necessary for long term olfactory habituation and promotes translation of the rate limiting circadian clock component, PERIOD (28, 29). In addition to TDP-43’s critical role in suppression of cryptic exons and splicing (8), we and others have shown that TDP-43 is actively transported with mRNA in the dendrites (30, 31) and axon as a component of neuronal ribonucleoprotein (RNP) transport granules (7, 32, 33). A key unresolved issue, however, is whether Ataxin-2 and TDP-43 interact functionally to regulate mRNA localization, stability, or translation.

In yeast and *Drosophila* models of TDP-43-induced neurodegeneration, the ability of Ataxin-2 to enhance TDP-43 toxicity requires the Ataxin-2 <u>p</u>oly(<u>A</u>)-binding protein-interacting (PAM2) motif and is RNA-dependent (15, 16). In mammalian cells, TDP-43 and Ataxin-2 co-immunoprecipitate in an RNA-dependent manner, and mutating phenylalanine residues in the TDP-43 RRM domains that are critical for RNA binding abrogates this biochemical interaction (16). Therefore, one possibility which has garnered support is that the genetic interplay between TDP-43 and Ataxin-2 polyQ intermediate-length expansions is mediated at the cellular level by enhanced recruitment of TDP-43 to cytoplasmic RNP granules, phase-separated condensates enriched in RNA and RNA-binding proteins (17, 34). In support of this notion, both TDP-43 and Ataxin-2 are components of stress granules, and ALS patient-derived lymphoblastoid cells possessing Ataxin-2 intermediate-length polyglutamine expansions show more robust cytoplasmic mislocalization of TDP-43 upon cellular stress compared to control cells (16, 35). Furthermore, detailed studies of the molecular determinants of condensate material properties predict that increased glutamine content may cause hardening and potentially alter condensate dynamics, but this has not been tested in neurons and the consequences for RNP condensate functions are not known (36, 37).

Using quantitative analysis of live cell and super-resolution imaging data, we and others have previously demonstrated that neuronal TDP-43 ribonucleoprotein granules display biophysical properties consistent with liquid-like condensates (33, 38). In this study, we show that wild type TDP-43 and Ataxin-2 exhibit distinct but overlapping spatial localization along the axon as well as striking differences in their dynamics and biophysical properties. Moreover, we hypothesized that Ataxin-2 polyQ expansions interact with TDP-43 at the cellular and molecular level to disrupt TDP-43 RNP condensate dynamics and post-transcriptional RNA regulation in neurons. Here we show that Ataxin-2 polyglutamine expansions not only perturb the liquid-like properties of TDP-43 RNP condensates, but also disrupt their anterograde transport and sequester TDP-43 in the proximal axon, leading to functional defects in mRNA localization and translation in neurons. Thus our findings provide insights into the contribution of Ataxin-2 polyQ expansion on TDP-43 dysfunction in ALS and offer a possible explanation for why motor neurons, with extremely long axons, are preferentially affected by Ataxin-2 polyQ expansions.

## RESULTS

### Spatial distribution and colocalization of TDP-43 with wild type Ataxin-2 in neuronal RNP condensates

TDP-43 localizes primarily within the nucleus, but also shuttles between the nucleus and cytoplasm in mammalian cell lines and fibroblasts (39). Similarly, neuronal TDP-43 is predominantly nuclear but is also distributed along the axon and dendrites in the form of RNP granules (7, 30-33). Using super-resolution microscopy of endogenous TDP-43 in primary cultured neurons, we and others have confirmed that TDP-43 forms rounded puncta, consistent with a de-mixed distribution, in both the nucleus and cytoplasm, including in the axon (33, 38). In contrast, Ataxin-2 is localized to the cytoplasm, with a tubulo-granular expression pattern in the soma and neurites, and associates with rough endoplasmic reticulum (40). However, colocalization between endogenous TDP-43 and Ataxin-2 has not been examined in neurons.

Considering that both TDP-43 and Ataxin-2 are components of RNP biomolecular condensates, such as stress granules (16, 29, 35, 41, 42), we first analyzed the expression and localization of endogenous TDP-43 and Ataxin-2 in rodent primary cortical neurons, DIV10-12. In agreement with prior studies, TDP-43 is predominantly nuclear with punctate expression in the dendrites and axon, whereas Ataxin-2 is localized in the soma as well as puncta in the dendrites and axon (Supplementary Fig. 1A-C; Fig. 1B,C). Consistent with their known functions in regulating mRNA stability and translation, we find that endogenous TDP-43 and Ataxin-2 co-localize with polyA mRNA as well as established markers of neuronal RNP granules, such as FMRP and PABPC (Fig. 1B-D, Supplementary Fig. 1D,E) (43, 44). We used a stringent objects-based colocalization analysis (45) to show that 27 ± 2.0% of TDP-43 granules and 26 ± 2.2% Ataxin-2 granules in the axon each colocalize with FMRP, a marker of neuronal transport granules (46). TDP-43 and Ataxin-2 colocalize to a significantly lesser extent with the canonical stress granule marker G3BP1 (p *β* 0.0003; Fig. 1D). Next, we examined the colocalization of endogenous TDP-43 and Ataxin-2 in the axon. We found that 23 ± 1.8% of TDP-43 granules colocalize with Ataxin-2, and a similar percentage of Ataxin-2 granules colocalize with TDP-43 (24 ± 1.9%) in the proximal axon (defined as <50 *μ*m from the soma) (Fig. 1C,E). Significantly fewer TDP-43 and Ataxin-2 double-positive granules are observed in the mid axon (p < 0.002; Fig. 1E). A histogram of the distribution of Ataxin-2 or TDP-43 along the axon shows that >93% of Ataxin-2 granules are localized in the proximal axon, whereas those positive for TDP-43 are distributed more evenly throughout the axon (Fig. 1F). These data indicate that TDP-43 and Ataxin-2 exhibit distinct but overlapping spatial localization along the axon and that their association with each other occurs predominantly in the proximal axon. Furthermore, their colocalization with FMRP and other known neuronal RNP markers suggest possible roles in co-regulation of mRNA transport and translation.

### RNP condensates positive for TDP-43 and/or Ataxin-2 show distinct motility and fluorescence recovery dynamics in the axon

We and others have previously shown that neuronal RNP condensates in different locations within the axon or dendritic compartments display distinct motility, biophysical, and functional properties (33, 47). Moreover, differences in RNA-binding protein composition affect RNP granule translocation from the dendritic shaft to spines, suggesting that RNA-binding proteins are important determinants of mRNA transport and localization (47). Therefore, we hypothesized that neuronal RNP condensates positive for TDP-43, Ataxin-2, or both proteins may show differences in their transport along the axon. We co-expressed fluorescently tagged human wild-type eGFP-TDP-43 and mScarlet-Ataxin-2 Q22, or either construct in combination with a control plasmid (e.g. fluorescently-tagged LAMP1) in rat primary cortical neurons (Supplementary Fig. 2). Exogenous expression of eGFP-TDP-43 WT and/or mScarlet-Ataxin-2 Q22 closely mirrors their endogenous expression pattern (compare Supplementary Fig. 1A-C, Supplementary Fig. 2A-D; (33)). We performed live-cell imaging to study the motility of neuronal RNP condensates positive for either TDP-43 or Ataxin-2 Q22, and those condensates positive for both TDP-43 and Ataxin-2 Q22 [TDP-43 (+) Ataxin-2 (+)] (Fig. 2A; Supplementary Fig. 2E,F). We found that the majority of TDP-43 positive condensates show either oscillatory (42 ± 5.0%) or long-range motility of ≥10 *μ*m net distance (20 ± 2.3%), whereas Ataxin-2 Q22 positive RNP condensates are predominantly stationary (95 ± 2.0%) (Fig. 2A,B). A histogram of the cumulative distance traveled by Ataxin-2 or TDP-43 positive condensates shows that only 1.4% of Ataxin-2 Q22 positive granules are transported ≥ 10 *μ*m while nearly 50% of TDP-43 granules travel cumulative distances of ≥ 10 *μ*m along the axon (Fig. 2C). Strikingly, TDP-43 (+) Ataxin-2 (+) condensates display motility similar to that of Ataxin-2 Q22 and are also predominantly stationary (80 ± 4.1%); only 6.3 ± 2.2% of double-positive condensates show long-range net transport ≥ 10 *μ*m (Fig. 2A,B). These data indicate that TDP-43 (+) Ataxin-2 (+) RNP condensates exhibit distinct transport compared to those that are TDP-43 (+) Ataxin-2 (-), suggesting that Ataxin-2 governs the motility of RNP condensates containing both Ataxin-2 and TDP-43.

**Figure 2.**
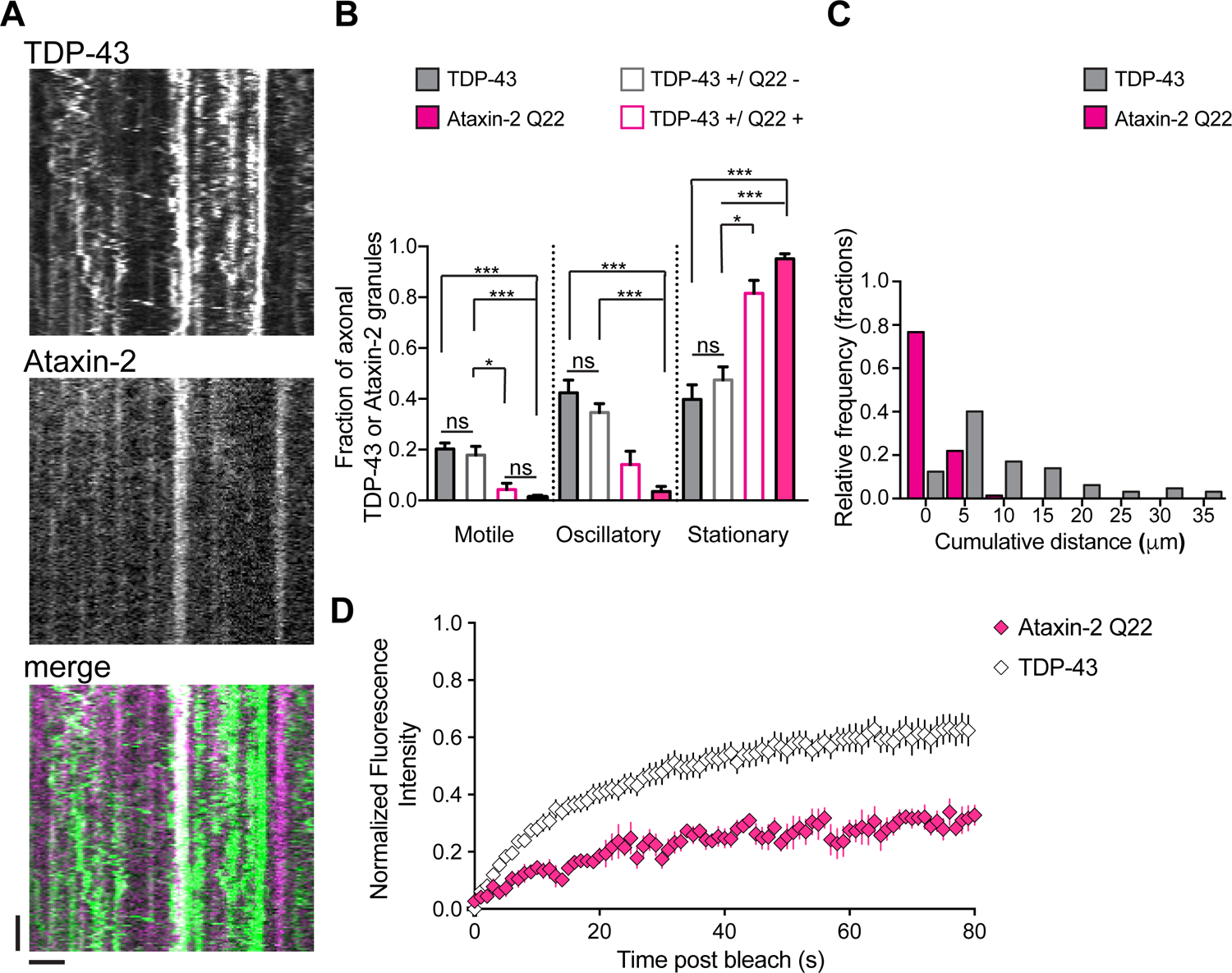
RNP condensates positive for TDP-43 and/or Ataxin-2 (Q22, wild type) show distinct motility and fluorescence recovery dynamics. **(A)** Representative kymographs of eGFP-TDP-43 and mScarlet-Ataxin-2 Q22 expressed in primary cortical neurons (DIV7-10); scale bar = 2.5 μm and 40s. **(B)** Kymographs were analyzed using custom MATLAB programs to quantify the motility of RNP condensates composed of TDP-43, Ataxin-2 Q22, or both TDP-43 and Ataxin-2 (TDP-43+/Q22+) in primary cortical neurons co-expressing eGFP-TDP-43 and mScarlet-Ataxin-2 Q22 (or control plasmids). The majority of RNP condensates containing TDP-43 without Ataxin-2 are motile (≥ 10 μm net distance) or oscillatory (< 10 μm net distance but > 5 μm cumulative distance) whereas Ataxin-2 positive condensates are predominantly stationary. Condensates positive for TDP-43 and Ataxin-2 are also predominantly stationary (*p<0.05, ***p<0.0005, Kruskal-Wallis test; n=16-21 neurons per condition, *N = 3*). **(C)** RNP condensates composed of eGFP-TDP-43 are transported over greater cumulative distances along the axon, compared to condensates composed of mScarlet-Ataxin-2 Q22 (p < 0.0001, Kolmogorov-Smirnov test; Ataxin-2 n=73, TDP-43 n=65 from 13-16 neurons each, *N = 3*). **(D)** Fluorescence recovery after photobleaching shows that Ataxin-2 displays slower and incomplete fluorescence recovery (*τ* = 25.5 s; A = 0.28) compared to TDP-43 (*τ* = 21.1 s; A = 0.58), suggesting that Ataxin-2 is a more stable component of RNP condensates (Ataxin-2 n=9, TDP-43 n=20, *N* = 3).

In addition to having different transport dynamics, next we asked whether Ataxin-2 also shows distinct fluorescence recovery dynamics compared to TDP-43. Using fluorescence recovery after photobleaching (FRAP), we observed that Ataxin-2 exhibits slower fluorescence recovery and a more limited mobile fraction after whole bleach compared to TDP-43 (Fig. 2D). Fluorescence recovery curves of Ataxin-2 and TDP-43 whole bleach analysis were fit to single exponential equations, in which A is the mobile fraction and *τ* = time constant (Table). TDP-43 displays more robust recovery (*τ* = 21.1 s; A = 0.58), suggesting that > 50% of TDP-43 exists in dynamic equilibrium with the non-bleached pool, whereas Ataxin-2 only recovers to 28% of its pre-bleach fluorescence intensity (*τ* = 25.5 s; A = 0.28). The distinct FRAP dynamics of TDP-43 and Ataxin-2 suggest that Ataxin-2 is a more stable, scaffolding component of neuronal RNP condensates (48). Taken together, the motility and fluorescence recovery data demonstrate that TDP-43 and Ataxin-2 have distinct dynamic properties and raise the possibility that condensates containing Ataxin-2 also may have distinct functional properties than those containing TDP-43 without Ataxin-2.

### Actively translating mRNA frequently colocalizes with Ataxin-2, but not TDP-43

Recent evidence suggests FMRP and TDP-43 co-regulate and repress translation, though only a few targets, including *Rac1* and *futsch/Map1B*, have been identified to date (31, 49-52). In contrast, Ataxin-2 co-sediments with polyribosomes in yeast and mammalian cells and enhances mRNA stability and translation of specific transcripts through PABPC/PAM2 dependent and independent mechanisms (26, 27). TDP-43 and Ataxin-2 co-immunoprecipitate in an RNA-dependent manner, suggesting that association with mRNAs may favor their assembly into RNP condensates (16). Crosslinking and immunoprecipitation (CLIP) studies have revealed that Ataxin-2 binds mRNAs via 3’UTR AU- and U-rich elements, whereas TDP-43 binds UG-rich elements (5, 6, 26). Of note, both TDP-43 and Ataxin-2 bind *tardbp* and *actb* (*β*-actin) mRNA, suggesting these RNA-binding proteins may co-regulate a handful of specific mRNA targets (26, 53). However, whether TDP-43 and Ataxin-2 differentially regulate the translation of shared mRNA targets is unclear.

Experimental approaches using live translation reporters suggest that mRNAs undergoing active translation show different subcellular localization and/or exhibit reduced dynamics compared to translationally repressed mRNAs, though this may not be the case for all transcripts (54, 55). Therefore, we hypothesized that the distinct spatial distribution and motility of RNP condensates positive for Ataxin-2, TDP-43, or both [TDP-43 (+) Ataxin-2(+)] may reflect differences in their translation states. We first used a variant of puromycin containing an alkyne group, O-propargyl-puromycin (OPP) (56), to measure synthesis of nascent polypeptides in wild type primary cortical neurons. Puromycin is a tRNA analog that causes premature chain termination during translation by entering the A site and results in the formation of a puromycinylated nascent chain (57, 58). OPP (± cyclohexamide) labeling, followed by Click reaction and immunofluorescence for endogenous Ataxin-2 and TDP-43 was used to visualize nascent polypeptide chains in relation to RNP granules containing these RNA-binding proteins (Supplementary Fig. 3A). We observed that TDP-43 rarely colocalizes with puromycin signal, whereas nearly a quarter of RNP granules containing Ataxin-2 colocalize with puromycin (Supplementary Fig. 3B). It has recently been shown, however, that a limitation of the puromycinylation assay is that the nascent peptide chain is released from the ribosome and diffuses away from the site of incorporation (59).

In order to examine the relationship between RNP condensates containing Ataxin-2 and/or TDP-43 and active translation sites with subcellular precision, we used the SunTag reporter assay to visualize the formation of nascent polypeptide chains and measure active sites of translation. Multiple scFV-sfGFPs bind to each nascent SunTag reporter protein, making the newly synthesized peptide visible above background fluorescence (54, 59, 60). In primary cortical neurons co-expressing the *β*-actin SunTag reporter, scFV-sfGFP, and fluorescently-tagged TDP-43 or Ataxin-2 Q22, we performed single molecule fluorescence in situ hybridization (smFISH) to visualize mRNA as well as sfGFP-positive nascent peptide (Fig. 3A). FISH-Quant, a MATLAB based image-analysis tool was used to quantify active sites of translation, defined by the colocalization of sfGFP reporter and mRNA signals, as previously described [Fig. 3A,B; (54, 61)]. Puromycin treatment served as a negative control and was able to effectively abolish sfGFP-positive sites of active translation (Fig. 3B; Supplementary Fig. 3C). Consistent with previous studies suggesting TDP-43 represses translation (50-52), the fraction of translating mRNA is mildly reduced overall in neurons expressing TDP-43 compared to those expressing Ataxin-2 (Supplementary Fig 3D). Moreover, active sites of translation were analyzed for co-localization with Ataxin-2 or TDP-43 to determine the fraction of translating mRNA associated with either RNA-binding protein (Fig. 3C-E). We found that 80 ± 4% of actively translating SunTag *β*-actin reporter mRNA colocalizes with RNP granules containing Ataxin-2 Q22. In contrast, only 29 ± 6% of actively translating reporter mRNA colocalizes with TDP-43 positive RNP granules (Fig. 3C-E). Taken together, our puromycinylation and SunTag data demonstrate a robust association between Ataxin-2 and actively translating mRNA, suggesting that the distinct spatial localization, motility, and biophysical properties of RNP condensates containing Ataxin-2 and TDP-43 may reflect distinct functions in translation regulation.

**Figure 3.**
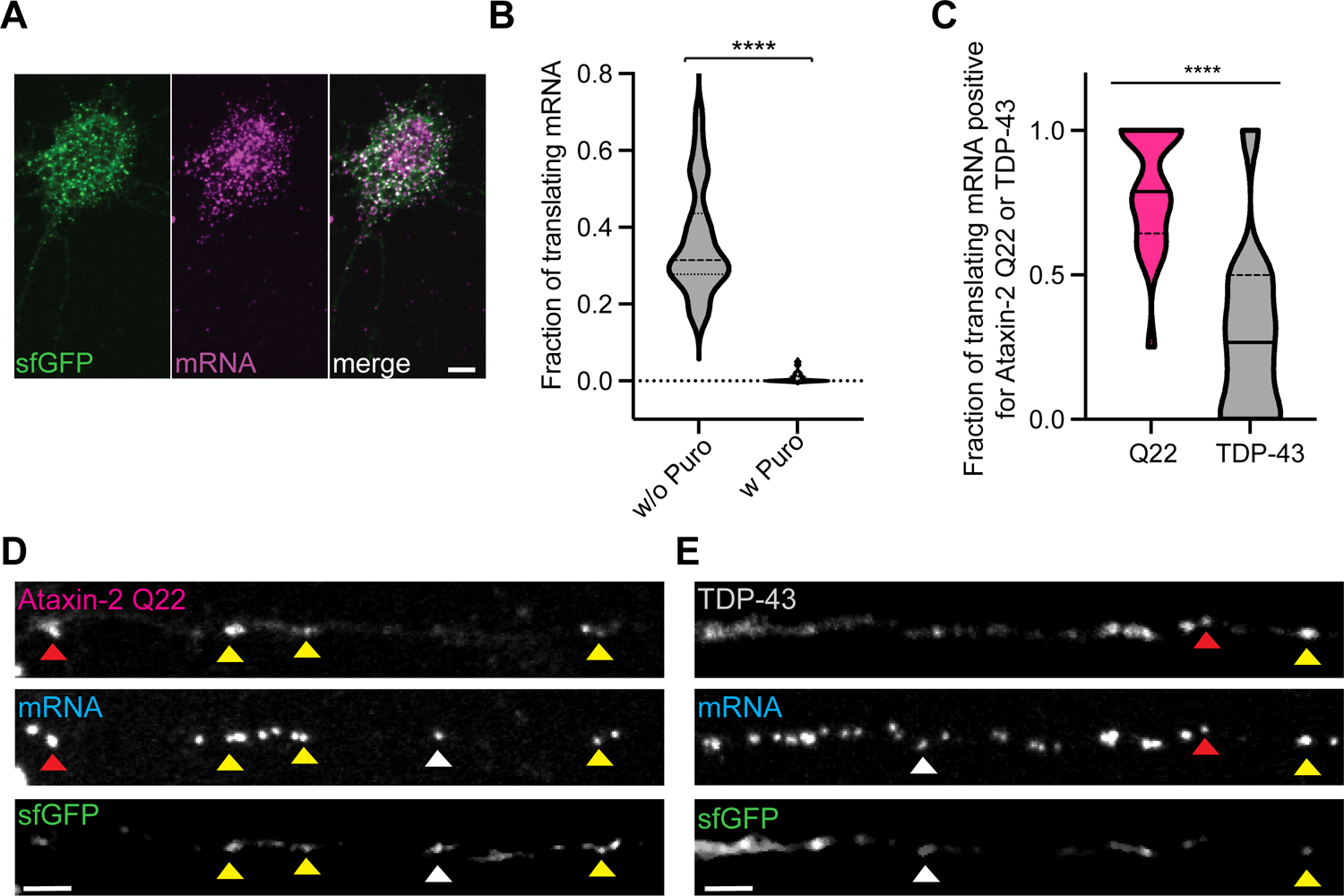
Actively translating SunTag reporter mRNA shows robust colocalization with Ataxin-2 positive RNP condensates. **(A)** Representative confocal image of SunTag *β*-actin translation reporter in a primary cortical neuron after fixation and smFISH to visualize SunTag mRNA. Green, sfGFP SunTag signal; magenta, mRNA; and white in the merged image indicates overlap between channels. Active sites of translation are defined by colocalization of mRNA with bright green sfGFP SunTag punctate signal (white spots in the merged image); scale bar = 5 μm **(B)** Fraction of translating mRNA decreases upon treatment with puromycin (100μg/mL, 90 min), (****p<0.0001, Mann-Whitney test; n=25 neurons per condition, *N=3*). (**C**) Fraction of translating mRNA that colocalizes with Ataxin-2 is significantly greater than the fraction of translating mRNA colocalizing with TDP-43 in dendrites (****p<0.0001, Mann-Whitney test; n=24 neurons per condition, *N=3*). **(D**,**E)** Representative confocal images of *β*-actin SunTag reporter mRNA, sfGFP, and either mScarlet-Ataxin-2 Q22 or mScarlet-TDP-43. Yellow arrowheads highlight actively translating mRNA that colocalize with Ataxin-2 **(D)** or TDP-43 **(E)**. Red arrowheads show RNP condensates positive for TDP-43 and mRNA, which are negative for sfGFP; scale bars = 3 μm.

### Ataxin-2 polyQ expansions disrupt axonal transport and sequester TDP-43

Intermediate length expansions of the Ataxin-2 polyglutamine tract are associated with increased risk for ALS, but the underlying cellular and molecular pathogenic mechanisms are unclear (16). TDP-43 regulates the stability and transport of synaptic and cytoskeletal transcripts, including *futsch*/*Map1B, Nefl*, and *STMN2*, and maintains local translation in the axon of motor neurons (7, 50, 62-66). Considering that Ataxin-2 regulates the motility of Ataxin-2(+) TDP-43(+) condensates (Fig. 2A,B), we investigated whether polyglutamine expansions of Ataxin-2 affect TDP-43 RNP transport, potentially leading to functional defects in mRNA localization that could contribute to ALS pathogenesis. We co-expressed eGFP-tagged wild type TDP-43 with mScarlet-Ataxin-2 of different polyQ lengths in primary cortical neurons and performed live cell confocal imaging. We then quantified the fraction of stationary, motile and oscillatory double positive [TDP-43 (+) Ataxin-2 (+)] condensates as well as TDP-43 (+) Ataxin-2 (-) condensates in the axon (Fig 4A). Similar to the axonal transport of condensates positive for TDP-43 ± wild type Ataxin-2 (Fig. 2), we observed that TDP-43 motility is largely determined by the presence or absence of Ataxin-2 within the RNP condensates (Fig. 4A). Neither increasing the length of Ataxin-2 polyQ tract, or removing it (*Δ*Q), significantly alters the motile, oscillatory, or stationary fraction of either TDP-43 (+) Ataxin-2 (+) or TDP-43 (+) Ataxin-2 (-) condensates (Fig. 4A). Instead, we found that in neurons co-expressing eGFP-TDP-43 and mScarlet-Ataxin-2 Q30 or Ataxin-2 Q39, the percentage of TDP-43 (+) Ataxin-2 (+) double positive RNP granules (46 ±3%) is significantly greater than in neurons co-expressing TDP-43 and Ataxin-2 Q22 (31 ± 4%) (Fig. 4B).

**Figure 4.**
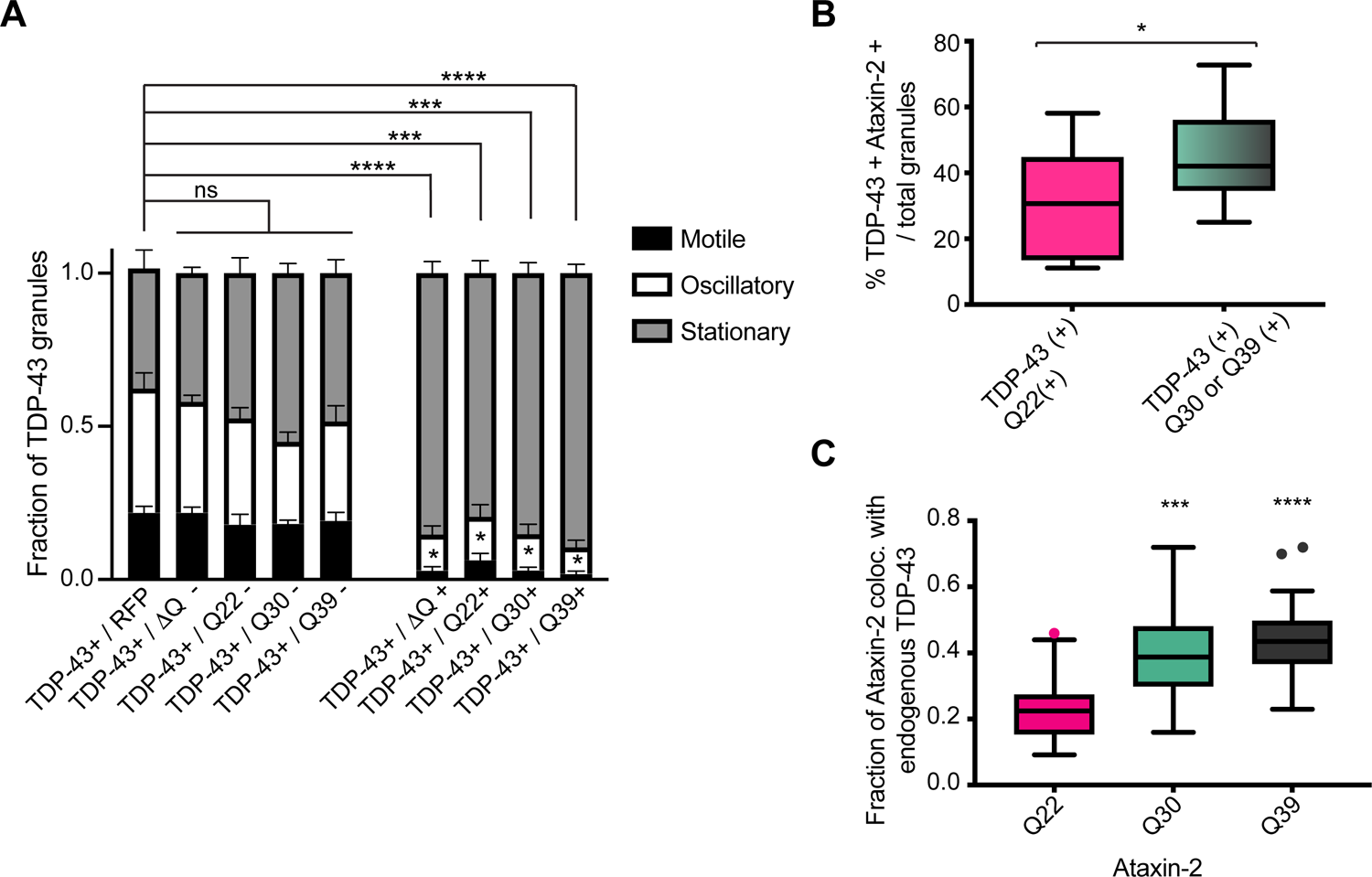
Ataxin-2 polyglutamine expansions disrupt motility and sequester TDP-43 in double positive TDP-43 (+) Ataxin-2 (+) condensates. **(A)** Fraction of motile, oscillatory, and stationary TDP-43 (+) RNP condensates positive or negative for Ataxin-2 polyQ variants. Regardless of Ataxin-2 polyQ length, TDP-43 (+) Ataxin-2 (+) condensates are predominantly stationary, compared to TDP-43 (+) Ataxin-2 (-) condensates (***p<0.0005; ****p<0.0001, Kruskal-Wallis test; n=15-21 neurons analyzed for each condition, *N=3*). **(B**) In neurons co-expressing eGFP-TDP-43 and mScarlet-Ataxin-2 Q30 or Ataxin-2 Q39, RNP condensates double positive for TDP-43 and Ataxin-2 account for a higher percentage of TDP-43 RNP granules compared to neurons expressing eGFP-TDP-43 and mScarlet-Ataxin-2 Q22 RNP (*p<0.02, Student’s t-test; n=14-17 neurons, *N=3*). **(C)** The fraction of Ataxin-2 colocalizing with endogenous TDP-43 is increased in the presence of Ataxin-2 polyQ expansions (***p<0.0005, ****p<0.0001, Kruskal Wallis test; n=22-25 neurons per condition, *N=3*).

Ataxin-2 and TDP-43 colocalize in abnormal cytoplasmic aggregates in ALS/FTD patient neurons (16). Therefore, we also examined whether Ataxin-2 polyQ expansions sequester endogenous TDP-43. Consistent with patient data, the fraction of Ataxin-2 positive RNP granules colocalizing with endogenous TDP-43 is also significantly increased in the presence of Ataxin-2 Q30 (37.0 ± 2.59%) or Ataxin-2 Q39 (44.1 ± 2.48%), compared to Ataxin-2 Q22 (23.6 ± 1.92%) (Fig. 4C). Similarly, we found that endogenous TDP-43 mislocalizes to Ataxin-2 Q30 and Q39 cytoplasmic aggregates in HEK293T cells, even in the absence of oxidative stress (Supplementary Fig. 4). In contrast, overexpression of fluorescently-tagged polyglutamine tract of similar length (BFP-Q30 and BFP-Q39), does not significantly alter colocalization of endogenous TDP-43 and Ataxin-2 (Supplementary Fig. 5A), suggesting that Q30 and Q39 polyglutamine tracts alone are not sufficient to aberrantly recruit endogenous TDP-43 to cytoplasmic condensates or aggregates. Taken together, these data suggest that Ataxin-2 polyQ expansions sequester TDP-43 within double positive condensates, most of which are stationary (Fig. 4A). Moreover, although the stationary fraction of TDP-43 is not increased by Ataxin-2 polyQ expansions, a higher fraction of TDP-43 is entrapped within those stationary TDP-43 (+) Ataxin-2 (+) condensates, potentially disrupting axonal transport of TDP-43 and crucial functions in the distal axon.

**Figure 5.**
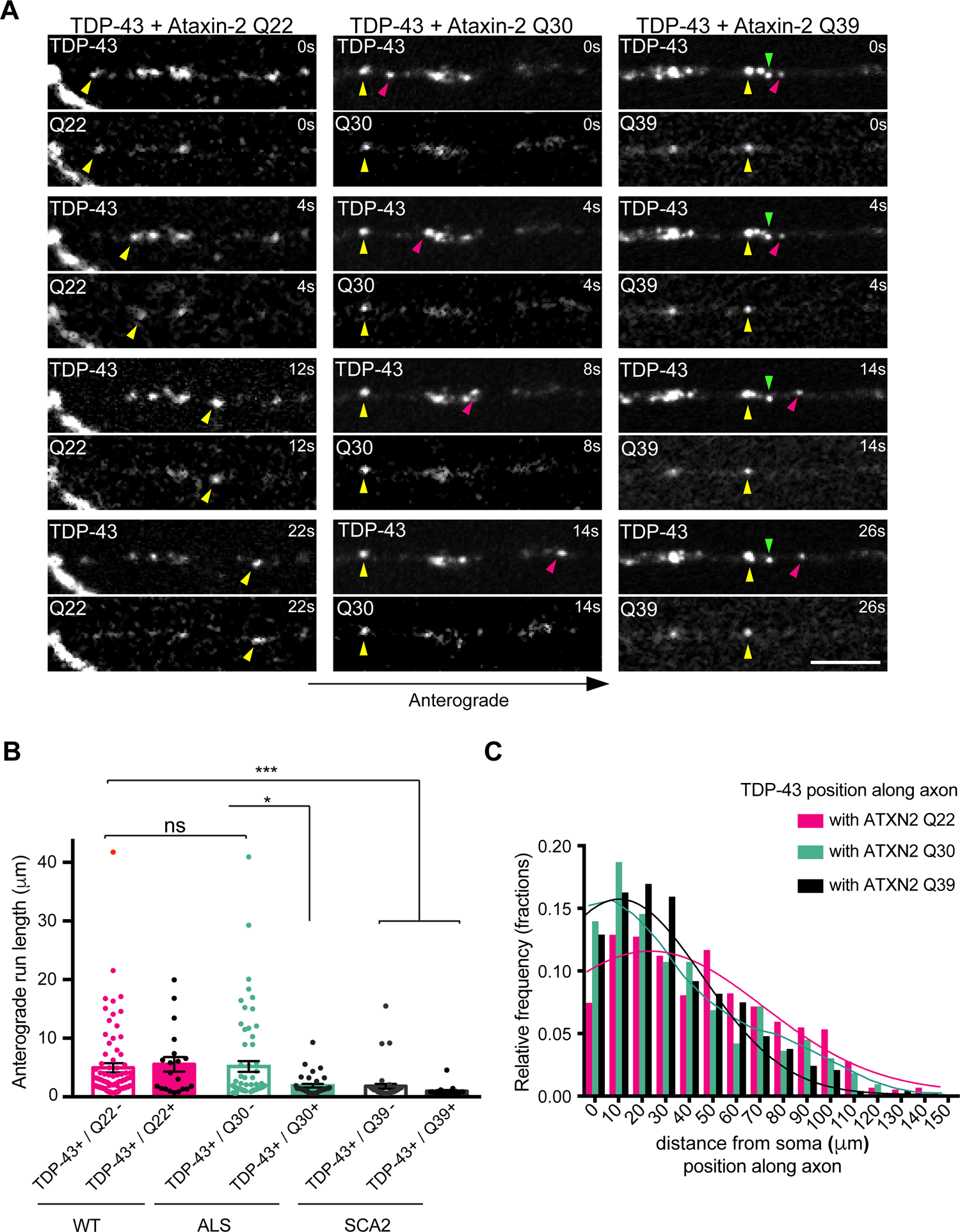
TDP-43 condensates positive for intermediate length Ataxin-2 polyQ expansions exhibit disrupted axonal anterograde motility. **(A)** Time-lapse imaging of eGFP-TDP-43 condensates in axons of primary cortical neurons expressing mScarlet-Ataxin-2 Q22, Ataxin-2 Q30, or Ataxin-2 Q39. Yellow arrowheads denote representative RNP condensates double positive for TDP-43 and Ataxin-2. TDP-43 (+) Ataxin-2 Q22 (+) condensates exhibit processive anterograde motility (yellow arrowhead, left). TDP-43 (+) Ataxin-2 Q30 (+) condensates show reduced anterograde run lengths (yellow arrowhead, middle), whereas run lengths of TDP-43 (+) Ataxin-2 Q30 (-) condensates (pink arrowhead, middle) are preserved. TDP-43 (+) Ataxin-2 Q39 (+) (yellow arrowhead, right) and TDP-43 (+) Ataxin-2 Q39 (-) condensates show limited anterograde movement (pink arrowhead, right) or are stationary (green arrowhead, right); Scale bar =3μm. **(B)** Anterograde run length quantitation of motile and oscillatory TDP-43 (+) RNP condensates positive or negative for Ataxin-2 Q22, Q30, or Q39 (*p<0.05; ***p<0.005, Kruskal-Wallis test, n=20-97 runs per condition from n=11-17 neurons, *N=3*). **(C)** Histogram of the position of TDP-43 condensates along the axon binned by distance (*μ*m) from the soma, in neurons expressing Ataxin-2 Q22, Q30, or Q39 (Kolmogorov-Smirnov, n=12 neurons per condition, *N=3*).

Next, we asked whether Ataxin-2 polyQ expansions inhibit long range processive transport of TDP-43 along the axon. Quantitative analysis of kymographs prepared from timelapse movies of primary cortical neurons co-expressing eGFP-TDP-43 and each mScarlet-Ataxin-2 polyQ variant showed that TDP-43 condensates containing Ataxin-2 Q30 or Ataxin-2 Q39 exhibit reduced anterograde run lengths, compared to those containing wild type Ataxin-2 Q22 (Fig. 5A, B; Supplementary movie). In neurons co-expressing Ataxin-2 Q22 and eGFP-TDP-43, there is no difference in anterograde run lengths of TDP-43 condensates, irrespective of their colocalization with Ataxin-2 Q22 [TDP-43 (+) Ataxin-2 Q22 (+) mean anterograde run length: 5.5 ± 1.2 *μ*m; TDP-43 (+) Ataxin-2 Q22 (-): 5.0 ± 0.8 *μ*m; Fig. 5B]. In contrast, TDP-43 condensates positive for Ataxin-2 Q30 show significantly lower anterograde run lengths (1.9 ± 0.3 *μ*m) compared to those that are negative for Ataxin-2 Q30 (5.2 ± 0.9 *μ*m; Fig. 5A, B). Unexpectedly, TDP-43 retrograde run lengths are similar in neurons expressing Ataxin-2 Q22 or Q30, regardless of colocalization with Ataxin-2 (Supplementary Fig. 5B). In neurons expressing Ataxin-2 Q39, however, both anterograde and retrograde run lengths of TDP-43 condensates are reduced regardless of whether they are positive for Ataxin-2 Q39 (anterograde: 1.0 ± 0.2 *μ*m; retrograde: −1.9 ± 0.5 *μ*m) or not (anterograde: 2.4 ± 0.4 *μ*m; retrograde: −2.5 ± 0.4 *μ*m, Fig. 5A, B, Supplementary. Fig. 5B). These data suggest Ataxin-2 Q30 intermediate length expansions are unique, and selectively deter anterograde transport of TDP-43 (+) Ataxin-2 Q30 (+) condensates only, while longer polyQ expansions may compromise the overall health of neurons and impair TDP-43 anterograde and retrograde run lengths in a non-specific manner. Given the reduced anterograde run lengths of TDP-43 (+) Ataxin-2 Q30 (+) condensates, we predicted that the spatial distribution of TDP-43 along the axon may be perturbed in the presence of Ataxin-2 polyQ expansions. We examined the position of TDP-43 condensates in the axon and plotted a histogram of TDP-43’s distance from the soma when co-expressed with each Ataxin-2 polyQ variants. We found that 50% of TDP-43 condensates are located within 25*μ*m of the soma in neurons expressing Ataxin-2 Q30 or Q39. In comparison, only 35% of TDP-43 condensates are found within 25*μ*m of the soma in neurons expressing Ataxin-2 Q22 (Fig. 5C). These findings suggest TDP-43 condensates are trapped proximally by Ataxin-2 with expanded polyQ tracts (Fig. 5B,C).

### Ataxin-2 polyQ expansions disrupt liquid-like properties of TDP-43 RNP condensates

We and others previously have shown that TDP-43 RNP granules exhibit liquid-like properties of biomolecular condensates in mammalian cells and in neurons (33, 38, 67). Our current results suggest TDP-43 is sequestered in RNP condensates positive for Ataxin-2 Q30 or Q39 and may display reduced dynamics. To test this prediction, we carried out fluorescence recovery after photobleaching (FRAP) experiments on axonal RNP condensates double positive for TDP-43 and Ataxin-2 Q22, Q30 or Q39. We analyzed fluorescence recovery of TDP-43 after whole bleach to quantify dynamic exchange of TDP-43 with the non-bleached cytoplasmic pool or half bleach to examine mobility of TDP-43 within the condensate (Fig. 6A-C). TDP-43 fluorescence recovery curves were fit to single exponential equations, in which A is the mobile fraction (Table). After whole bleach (Fig. 6C, top), fluorescence recovery of TDP-43 within RNP condensates containing Ataxin-2 Q22 is robust (A = 0.63, Table), suggesting that ∼ 60% of TDP-43 exists in dynamic equilibrium with the non-bleached pool. The mobile fraction of TDP-43 is progressively reduced in RNP condensates containing Ataxin-2 Q30 (A = 0.46) and Ataxin-2 Q39 (A = 0.35) indicating that exchange with the cytoplasmic pool of non-bleached TDP-43 is restricted in the presence of Ataxin-2 polyQ expansions. Furthermore, half-bleach fluorescence recovery data demonstrate that mobility of TDP-43 within condensates positive for Ataxin-2 polyQ expansions is slower and markedly reduced (Ataxin-2 Q30: A = 0.36, *τ* = 17.7s; Ataxin-2 Q39: A = 0.30, *τ* = 33.1s) compared to those positive for Ataxin-2 Q22 (A = 0.51, *τ* = 13.6s; Table, Fig. 6C bottom panel). This disruption in dynamic exchange and liquid-like properties of TDP-43 when associated with Ataxin-2 polyQ expansions could contribute to TDP-43 loss of function in disease pathogenesis and impair transport and translation of critical mRNA transcripts in the distal axon. In addition, these findings raise the possibility that Ataxin-2 polyQ expansions may also sequester other RNA-binding proteins.

**Figure 6.**
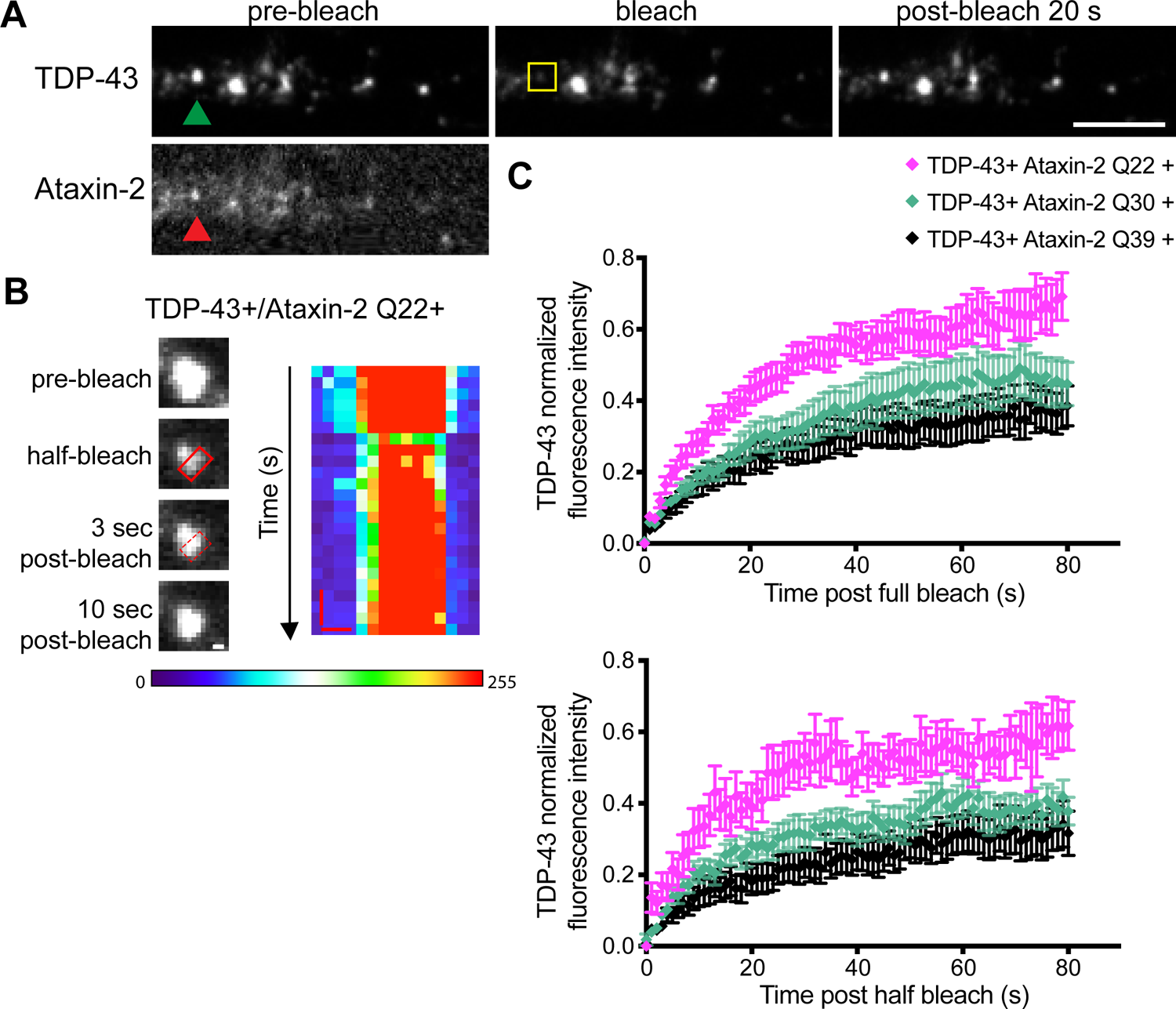
Fluorescence recovery after photobleaching experiments show that Ataxin-2 polyQ expansions impair TDP-43 RNP condensate dynamics. **(A, B)** RNP condensates double positive for TDP-43 (green arrowhead) and Ataxin-2 Q22 (red arrowhead) show robust recovery of TDP-43 fluorescence intensity after **(A)** whole-or **(B)** half-bleach, suggesting that dynamic exchange between condensates and the soluble pool of TDP-43 and reorganization within TDP-43 condensates occur. Rectangular boxes highlight the bleached area; in **(A)**, scale bar=5 μm and in **(B)** scale bar=0.3 μm. In the heat map **(B)**, red denotes high fluorescence intensity and blue represents background intensity; scale bars= 0.3 μm, 3 s). **(C)** Top: TDP-43 fluorescence recovery after whole bleach is impaired in neuronal RNP condensates positive for either Ataxin-2 Q30 or Ataxin-2 Q39, indicating reduced dynamic exchange with the non-bleached pool of TDP-43 (n=10-11 condensates per condition, *N=3*). Bottom: TDP-43 fluorescence recovery after half-bleach shows impaired molecular mobility of TDP-43 within condensates containing Ataxin-2 polyQ expansions, suggesting the liquid-like properties of TDP-43 condensates is also perturbed (n=8-10 condensates per condition, *N=3*).

Indeed Ataxin-2 CAG42 and CAG100 knock-in mouse models have shown that Ataxin-2 polyQ expansions in the SCA2 range sequester PABPC1, TDP-43 and potentially other splicing factors (68, 69). We focused on G3BP1, a stress granule marker and key regulator of axonal translation in neurons (70), which shows a lower degree of colocalization with endogenous TDP-43 and Ataxin-2 under physiologic conditions (Fig. 1D). Consistent with Ataxin-2 KI mouse *in vivo* studies, we found that in primary cortical neurons expressing Ataxin-2 Q30 or Ataxin-2 Q39, the fraction of Ataxin-2 RNPs colocalizing with endogenous G3BP1 is significantly greater (Ataxin-2 Q30: 52.8 ± 3.6 %; Ataxin-2 Q39: 53.5 ± 3.1%) than in neurons expressing Ataxin-2 Q22 or a control plasmid with fluorescently-tagged Q30 or Q39 tract (Ataxin-2 Q22: 23.9 ± 1.9%; Supplementary Fig. 5C). The fraction of endogenous TDP-43 colocalizing with G3BP1 is similarly increased in neurons expressing Ataxin-2 Q30 or Ataxin-2 Q39 (Supplementary Fig. 5D). Removing the Ataxin-2 polyQ tract (Ataxin-2 ΔQ) does not significantly alter colocalization between Ataxin-2 and G3BP1 or TDP-43 and G3BP1, either (Supplementary Fig. 5C,D). Taken together, these data suggest Ataxin-2 polyQ expansions aberrantly sequester TDP-43 and possibly other RNA-binding proteins, and thereby may affect RNP condensate composition, dynamics, and functions in mRNA localization and translation.

### Ataxin-2 polyQ expansions disrupt mRNA localization in the mid axon

Since TDP-43 regulates mRNA stability, transport, and localized protein synthesis (65), which are important regulators of axonal health, we investigated whether Ataxin-2 polyQ expansions affect mRNA localization in neurons. *β*-actin mRNA is one of the most abundant mRNAs in neurons, directly binds with Ataxin-2 (26) and is transported by TDP-43 RNP granules (53, 71). To examine functional consequences of Ataxin-2 polyQ expansions on mRNA localization, we performed smFISH for *β*-actin mRNA on primary cortical neurons expressing mScarlet-tagged Ataxin-2 Q22, Ataxin-2 Q30, Ataxin-2 Q39 or a control plasmid and quantified the fluorescence intensity of mRNA in the soma and in mid to distal regions of the axon (Fig. 7A). There are no significant differences in the fluorescence intensity of *β*-actin mRNA in the soma for any of these conditions (Fig. 7B). However, neurons expressing Ataxin-2 Q30 or Ataxin-2 Q39, show reduced fluorescence intensity of *β*-actin mRNA in the axon compared to controls (Fig. 7A, C), suggesting reduced stability and/or long range axonal transport of this transcript. These results suggest polyQ expansions not only affect RNP condensate dynamics, but also are linked to functional defects in localization of axonal *β*-actin mRNA, and possibly other essential transcripts, and may contribute to ALS pathogenesis.

**Figure 7.**
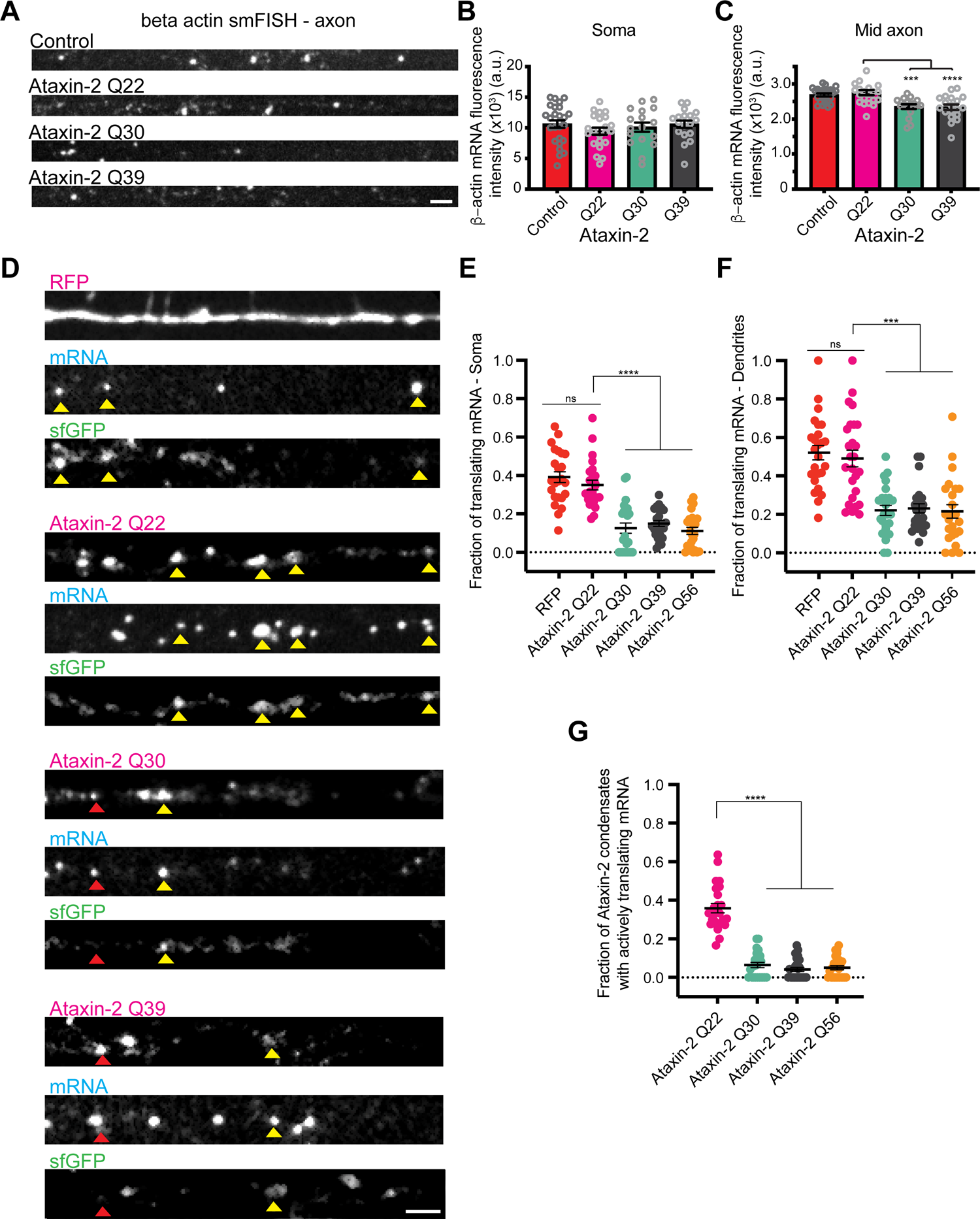
Ataxin-2 polyglutamine expansions disrupt mRNA localization and local translation in neurons. **(A)** Representative confocal images of smFISH for endogenous β-actin mRNA in mouse primary cortical neurons (DIV 9) transfected with either control (RFP) or mScarlet-Ataxin-2 Q22, Ataxin-2 Q30, or Ataxin-2 Q39; scale bar = 2.5*μ*m. **(B, C)** Quantification of β-actin mRNA fluorescence intensity **(B)** in the soma and **(C)** in the mid axon (50-150 *μ*m away from the soma) shows reduced fluorescence intensity in the axon of neurons expressing Ataxin-2 Q30 or Ataxin-2 Q39, compared to control or Ataxin-2 Q22. (***p<0.0005, ****p<0.0001, Kruskal_Wallis test; n=24 neurons per condition, *N=3*). **(D)** SunTag *β*-actin translation reporter was used to visualize active sites of translation in primary cortical neurons expressing reporter plasmids and control (RFP) plasmid or mScarlet-tagged Ataxin-2 polyQ variants. Images of dendrites in which mRNA colocalizing with sfGFP (yellow arrowheads) indicate an active site of translation. Ataxin-2 Q22 frequently associates with local translation sites. In contrast, mRNA rarely associates with sfGFP in condensates positive for Ataxin-2 Q30 or Q39 (red arrowheads). Scale bar =3μm. **(E)** Fraction of translating mRNA is perturbed in neuronal soma expressing Ataxin-2 polyQ expansions. (****p<0.0001, Kruskal-Wallis test; n=24, *N=3*). **(F)** Fraction of translating mRNA is suppressed in dendrites of neurons expressing Ataxin-2 polyQ expansions. (****p<0.0001, Kruskal-Wallis test; n=25, *N=3*). **(G)** Fraction of Ataxin-2 associating with active translation sites is significantly reduced in neurons expressing Ataxin-2 polyQ expansions. (****p<0.0001, Kruskal-Wallis test; n=26, *N=3*).

### Local translation is suppressed in RNP condensates positive for Ataxin-2 polyQ expansions

In addition to regulating mRNA transport, TDP-43 also controls many other aspects of RNA metabolism, including translation of ribosomal protein mRNAs, which may affect global and/or compartment specific translation (72). Phase separation defective TDP-43 causes alteration in global translation (73). Similarly, Ataxin-2 is reported to mediate post-transcriptional polyadenylation and enhance translation of its target mRNAs (26, 74), and promote global translation through its association with the translation pre-initiation complex (75). In non-neuronal cells, Ataxin-2 polyQ expansions (Q31-39) modestly reduce the expression of Ataxin-2 target mRNAs (26). Based on these studies and our own data (Fig. 3), we hypothesized that Ataxin-2 polyQ expansions may impair global translation and functionally impact RNP condensate regulation of local translation in neurons.

In order to determine whether translation is affected by Ataxin-2 polyQ expansions, we performed a puromycinylation assay with O-propargyl-puromycin (OPP) (56) on neurons expressing mScarlet-Ataxin-2 of differing polyQ lengths or a control plasmid (Supplementary Fig. 6). Pretreatment with cycloheximide suppresses puromycin incorporation into nascent polypeptides and thus served as a negative control (Supplementary Fig. 3A). In neurons expressing Ataxin-2 Q30 or longer polyQ tracts, we observed fewer signals for puromycinylated nascent chains along the axon (Ataxin-2 Q30: 0.06 ± 0.01 puromycin puncta/*μ*m) compared to neurons expressing Ataxin-2 Q22 or control (0.20 ± 0.02 puncta/*μ*m; Supplementary Fig 6A,B). This effect was more pronounced with longer polyQ tracts (Ataxin-2 Q39 and Ataxin-2 Q56). These findings suggest translation is suppressed in neurons expressing Ataxin-2 with longer polyQ tracts, but do not distinguish between global and local effects on translation. In addition, these data could also be consistent with defects in transport of newly synthesized proteins into the axon.

Next, we sought to examine whether Ataxin-2 polyQ expansions are associated with defects in local translation. Our data in Fig. 3 suggest that actively translating reporter mRNA shows frequent colocalization with condensates positive for Ataxin-2 Q22. Therefore, we hypothesized that neuronal RNP condensates containing Ataxin-2 with intermediate length or longer polyQ tracts will show reduced association with active translation sites. To address these questions, we again utilized the *β*-actin SunTag reporter assay in primary cortical neurons (54). We co-expressed *β*-actin SunTag reporter, scFV-sfGFP, and either a control plasmid, mScarlet-Ataxin-2 Q22, or its polyQ variants and performed smFISH to visualize the reporter mRNA. Importantly, there were no significant differences in the number of reporter mRNA puncta observed in neurons expressing Ataxin-2 Q22 or any of the Ataxin-2 polyglutamine expansions, indicating similar expression level of the SunTag reporter mRNA in all of the conditions tested (Supplementary Fig. 6C, D). We found that the fraction of translating *β*-actin mRNA is significantly decreased in neurons expressing Ataxin2 with polyQ expansions, compared to Ataxin-2 Q22 or control plasmid (Fig. 7D-F). We observed a greater than 50% reduction in the fraction of translating mRNA both in the soma and dendrites, suggesting translation is suppressed across multiple subcellular compartments. We also observed a low fraction of translating *β*-actin mRNA in the axon (not shown); although this sparse signal could not be analyzed for statistically robust differences between the control and Ataxin-2 polyQ variants, we expect axonal translation is also affected, as seen in the soma and dendrites.

Since intermediate length and longer Ataxin-2 polyQ expansions were associated with impaired translation in neurons, next we asked whether RNP condensates positive for expanded Ataxin-2 polyQ variants show evidence of translation dysregulation. In this analysis we quantified the number of actively translating *β*-actin mRNA that associates with Ataxin-2 (i.e., triple positive for mRNA, sfGFP, and Ataxin-2 variant) as a fraction of the total number of Ataxin-2 positive RNP condensates. We found that RNP condensates containing Ataxin-2 polyQ expansions rarely colocalize with active sites of translation (Fig. 7G). Only 6.4 ± 1.4 % and 4.2 ± 1.1% of RNP condensates positive for Ataxin-2 Q30 and Ataxin-2 Q39, respectively, colocalize with actively translating mRNA. In comparison, 36 ± 2.4% of RNP condensates positive for Ataxin-2 Q22 colocalize with actively translating reporter mRNA (Fig. 7G). These data suggest RNP condensates positive for Ataxin-2 polyQ expansions not only exhibit impaired dynamics and transport, but also are linked to functional defects in local translation. Moreover, these findings support and expand upon growing evidence that dysregulation of mRNA localization and translation have detrimental effects on neuronal integrity and contribute to ALS pathogenesis (65, 66).

## DISCUSSION

In this study, we examined the transport dynamics, biophysical properties, and mRNA regulatory functions of RNP condensates positive for TDP-43 and/or Ataxin-2 in neurons, large and morphologically complex cells. Our data reveal functional properties of neuronal RNP condensates that are novel and not evident in studies of condensates in cell lines. We show that RNP condensates containing TDP-43 or Ataxin-2 exhibit distinct but overlapping spatial localization along the axon, different transport dynamics and each display unique biophysical properties, all of which may contribute to their distinct roles in translation regulation. Moreover, our data for the first time reveal mechanisms by which Ataxin-2 polyglutamine tract expansions disrupt RNP condensate functions, specifically mRNA localization and regulation of local translation, by perturbing axonal transport and the dynamic liquid-like properties of TDP-43 as well as by altering RNP condensate composition (Fig. 8).

**Figure 8.**
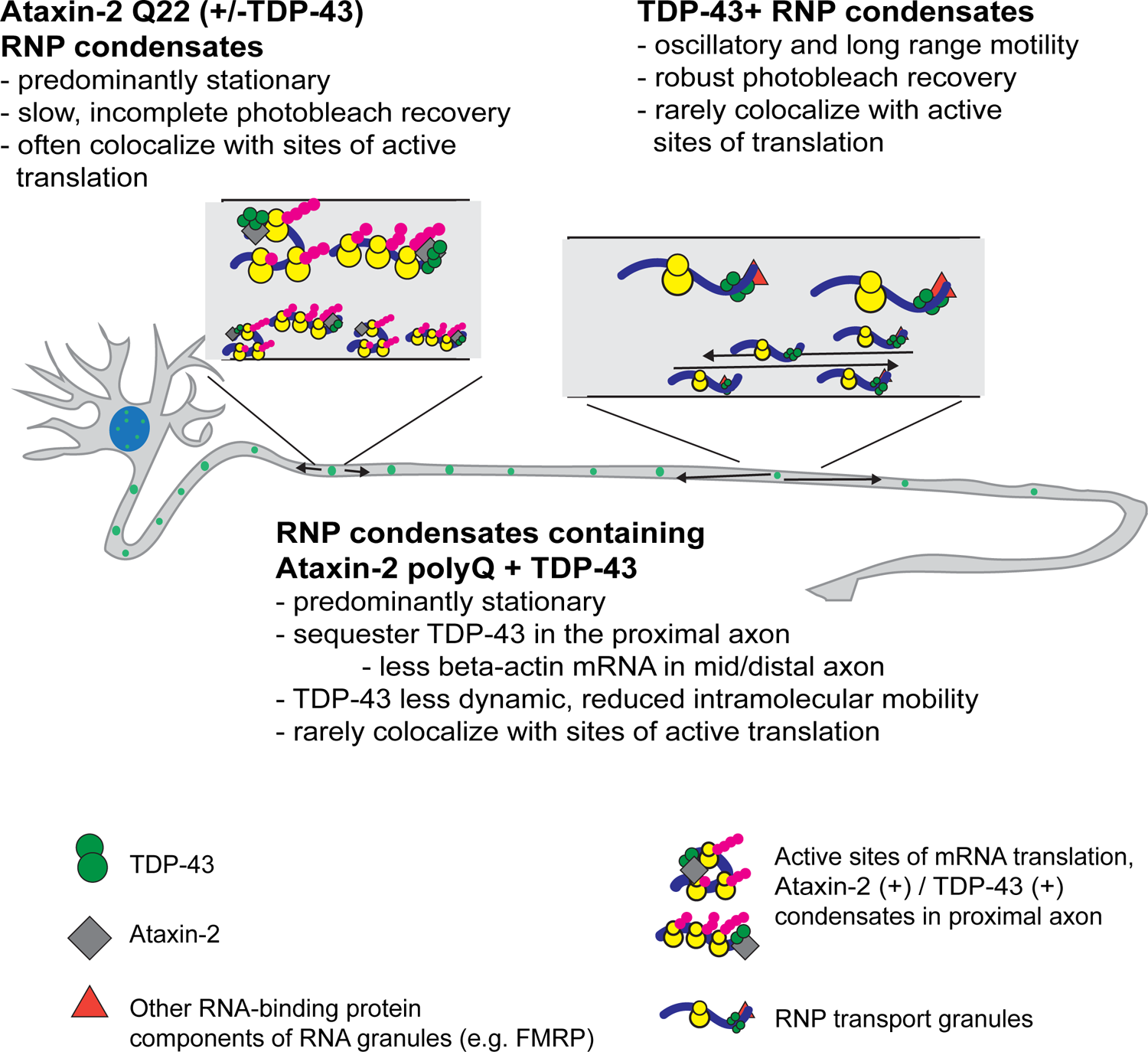
**Summary schematic** of localization, motility and functional properties of RNP condensates containing TDP-43 and wildtype Ataxin-2 or polyQ variants in neurons.

The size and morphologic complexity of neurons, including the length of the axon and dendritic arborization, impose logistical challenges to maintain synaptic proteins and organelle homeostasis at remote locations from the nucleus. Neurons rely heavily on cytoskeletal transport mechanisms in order to maintain protein, organelle and synaptic function (76). In addition, the discovery of thousands of mRNAs near synaptic sites suggest that neurons, in part, also rely on local protein synthesis (77, 78). FMRP, TDP-43 and other RNA-binding proteins form cytoplasmic RNP transport granules that facilitate delivery of target mRNA to distal neuronal compartments. RNA-binding protein composition is a critical determinant of mRNA transport (79, 80), and indeed, our data show that RNP condensates positive for TDP-43, Ataxin-2, or both [TDP-43(+) Ataxin-2(+)] display markedly different motility and localization in the axon. Whereas RNP granules positive for TDP-43 display oscillatory or long-range motility (net transport *β*10*μ*m), RNP condensates positive for Ataxin-2, with or without TDP-43, are largely stationary. Thus our data identify Ataxin-2 as an important determinant of motility in neuronal RNP condensates and highlight how a single RNA-binding protein can significantly impact RNP function.

Furthermore, we show that actively translating mRNA colocalizes with Ataxin-2 to a significantly greater degree than with TDP-43. Our findings are consistent with and expand upon the observation that Ataxin-2 associates with polyribosomes and rough endoplasmic reticulum (27, 40). In contrast, less than 30% of the translating *β*-actin mRNA colocalizes with TDP-43(+) RNP condensates. Whether the subset of TDP-43 (+) condensates that associate with translating mRNA also contain Ataxin-2 is unclear, however, since we were not able to simultaneously visualize more than one RNA-binding protein along with mRNA and nascent peptide. Future studies will be required to identify the RNA-binding proteins or other regulatory components in translationally active TDP-43(+) RNP condensates, and whether TDP-43 and Ataxin-2 play synergistic or opposing roles in regulating the translation of common target mRNAs. Taken together, our data bolster evidence that spatial localization, motility, and RNA-binding protein composition of neuronal RNP condensates are important factors in determining translation states of mRNA (55).

Prior studies suggest RNP granules in neurons are highly dynamic and that this is an important feature of their ability to deliver mRNAs to distal sites for local translation (81, 82). We found that the fluorescence recovery dynamics of wild type Ataxin-2 and TDP-43 within RNP condensates are also distinct, raising the possibility they play distinct functional roles. The robust fluorescence recovery dynamics of TDP-43 suggest that it may act as a client molecule, whereas the low mobile fraction and slow fluorescence recovery of Ataxin-2 would be more consistent with a role as a scaffold molecule within the condensate (48). Studies showing that knockdown of Ataxin-2 reduces stress granule formation supports its potential role as a scaffold (17), but confirmation of these distinct roles for TDP-43 and Ataxin-2 in neuronal RNP condensates awaits additional investigation.

ALS has been described as a distal axonopathy (83), and there is emerging evidence that defects in stability, localization, and local translation of transcripts critical for maintenance of the microtubule cytoskeleton, axon caliber, and oxidative phosphorylation are key features of ALS/FTD (51, 65, 66, 84). In addition to regulating mRNA splicing and stability, TDP-43 is important for transport of target mRNAs from the soma to distal axonal compartments, including the NMJ. ALS-linked TDP-43 mutations impair this mRNA localization function in stem cell-derived motor neurons from ALS patients (7). Here we used several approaches to show that Ataxin-2 polyQ expansions sequester TDP-43 and disrupt motility of TDP-43 condensates along the axon. TDP-43 RNP granules containing Ataxin-2 Q30 exhibit selective axonal anterograde motility defects, in contrast to granules that are negative for the intermediate-length Ataxin-2 polyQ variant. As a result, Ataxin-2 polyQ expansions lead to TDP-43 accumulation in the proximal axon, with potential functional defects at the NMJ. In support of this, we found that Ataxin-2 polyQ expansions decrease the localization of endogenous *β*-actin mRNA to the mid axon, suggesting that stability and/or transport of *β*-actin mRNA to distal regions is diminished. While *β*-actin mRNA transport is also regulated by other RNA-binding proteins, such as zipcode-binding protein 1 (ZBP1/IMP1/IGF2BP1) (85, 86), we speculate that there will be more profound defects in distal localization of mRNA targets regulated exclusively by TDP-43.

In addition, we found that Ataxin-2 polyQ expansions sequester TDP-43 and perturb its liquid-like properties within RNP condensates. Using FRAP assays, we showed that condensates positive for TDP-43 and Ataxin-2 Q30, or Ataxin-2 Q39, exhibit reduced dynamic exchange with the non-bleached pool of TDP-43 and slower intra-condensate mobility, compared to those containing TDP-43 and Ataxin-2 Q22. Consistent with the notion that Ataxin-2 polyQ expansions alter RNP condensate dynamics, in our study we also observed an increased fraction of double positive TDP-43 (+) Ataxin-2 (+) RNP granules in neurons expressing Ataxin-2 with expanded polyQ tracts, suggesting there is sequestration of endogenous TDP-43. Previous reports have shown that cytoplasmic TDP-43 liquid-liquid phase separation drives a feedforward mechanism compromising nucleo-cytoplasm transport and exacerbating loss of nuclear TDP-43 (38). Although we did not observe loss of nuclear TDP-43, mutant Huntingtin was shown previously to sequester TDP-43 and compromise splicing (87), and further studies are needed to investigate whether Ataxin-2 polyQ expansions impact nuclear TDP-43 splicing function. Altogether, these data suggest that Ataxin-2 polyQ expansions may trigger aberrant phase transitions and contribute to axonopathy through sequestration of TDP-43 and critical axonal transcripts within RNP condensates.

Recently, enhanced levels of phosphorylated TDP-43 were found in postmortem spinal cord tissue harboring intermediate length Ataxin-2 expansions (88). Phosphorylated TDP-43 binds and sequesters nuclear-encoded mitochondrial mRNAs as well as G3BP1, a negative modulator of intra-axonal protein translation, thus depleting essential mitochondrial proteins in the axon, which leads to neurodegeneration (65, 70). We found Ataxin-2 polyQ expansions similarly sequester TDP-43 as well as G3BP1 in neuronal RNP condensates, and that translation of reporter mRNA is markedly reduced across multiple cellular compartments. Notably, this effect is even more striking when we quantified the fraction of active translation sites that colocalize with Ataxin-2 Q22 as opposed to those with Ataxin-2 Q30 or Q30. Importantly, these results suggest that Ataxin-2 polyQ expansions not only perturb RNP condensate transport and liquid-like properties, but also have detrimental functional outcomes. Taken together, our findings as well as several recent studies indicate that misregulation of local translation, specifically of critical axonal mRNAs, could contribute to a “dying back” pattern of neurodegeneration.

In summary, we show that RNA-binding protein composition is an important determinant of spatial localization, transport dynamics, and translation of neuronal RNP condensates, and that Ataxin-2 plays a critical role in post-transcriptional RNA homeostasis. Moreover, this work for the first time reveals mechanisms underlying aberrant interactions between TDP-43 and Ataxin-2 polyQ expansions that have detrimental effects on transport, localization and liquid-like biophysical properties of TDP-43 and contribute to toxic gain-of-function consequences for local translation as well as mRNA localization. Overall, our findings provide a framework for why motor neurons with long axons may be particularly susceptible for neurodegeneration associated with Ataxin-2 polyglutamine expansions.

## ACKNOWLEDGEMENTS

We are grateful to Dr. Junjie Guo, Dr. Janghoo Lim (Department of Neuroscience, Yale University), and members of the Gopal lab for their thoughtful discussion and comments on the manuscript. We thank Daniel Calbick (Interdepartmental Neuroscience Program, Yale University) for creating a custom-written MATLAB pipeline to de-noise time-lapse microscopy images of axonal transport and Dr. Sabyasachi Sutradhar (Yale University) for his help with creating a custom MATLAB script to analyze co-localization of mRNA with nascent peptides. This research was supported by the ALS Association (Award ref. 19-IIA-487) and the National Institute of Neurological Disorders and Stroke/NIH under Awards K08-NS094744 and R01NS122907 (to PPG).

## AUTHOR CONTRIBUTIONS

DW and PPG conceptualized and designed the study, performed experiments, acquired data, analyzed and interpreted data and wrote the manuscript. NV and SSV aided in data acquisition and edited the manuscript. PPG supervised the entire study. All authors contributed to manuscript revision and approved the submitted version.

## MATERIALS AND METHODS

### Plasmids

Constructs include eGFP-TDP-43 WT (gift from Dr. Virginia Lee), mScarlet-Ataxin-2 Q22, Q30, Q39, Q56, or *Δ*Q (polyQ deleted). mScarlet-tagged Ataxin-2 constructs were generated from pcDNA6-Ataxin-2 Q22, Q30, and Q39 (gifts from Dr. Aaron Gitler). Ataxin-2 *β*Q and Ataxin-2 Q56 were generated using site directed mutagenesis PCR and the primers given in Table 1. To generate Q30 and Q39 control plasmids, the polyglutamine tracts of mScarlet-Ataxin-2 Q30 and Q39 were amplified using primers given in Table 1. Q30 and Q39 PCR products were subcloned into pJet 1.2 blunt cloning vector (ThermoFisher) prior to insertion into eBFP-C1 plasmid using Kpn I and Bgl II sites. Additional plasmids were obtained from Addgene: pUbC-OsTIR1-myc-IRES-scFV-sfGFP (Addgene plasmid # 84563) and pUbC-FLAG-24xSuntagV4-oxEBFP-AID-baUTR1-24xMS2V5-Wpre (Addgene plasmid # 84561) were gifts from Dr. Robert Singer.

### Primary cortical neuron culture and transfection

Primary cortical neurons were dissociated from brains of E18 Sprague-Dawley rat embryos or E16 C57Bl/6 mouse embryos and cultured in Neurobasal medium (Gibco) as previously described (Gopal *et al*., 2017). Neurons were plated at a density of 150,000 cells/mL on poly-L-lysine (0.5mg/mL; Sigma) coated coverslips or MatTek dishes in attachment media [MEM containing 10% horse serum (Gibco), 1mM pyruvic acid (Gibco) and 33mM glucose (Gibco)] for 2-4 hours. Neurons were maintained in Neurobasal media (Gibco) containing B27 supplement (Invitrogen), 2mM GlutaMAX (Gibco), 33mM glucose (Gibco), 100units/mL penicillin and 100*μ*g/mL streptomycin at 37 °C with 5% CO_2_. On the third day in vitro (DIV 3), 1*μ*M AraC was added. For live cell imaging, rat primary cortical neurons (DIV 7-9) were co-transfected with eGFP-TDP-43 WT and mScarlet-Ataxin-2 polyQ variants or control plasmid (total DNA =1.5*μ*g) using Lipofectamine 2000 (Thermo Fisher Scientific).

### Mammalian cell culture and transfection

HEK293T (ATCC) cells were cultured and maintained in DMEM, 10% FBS and 1% 100X GlutaMAX. For stress granule experiments, HEK293Tells were transfected with ATAXIN-2 Q22, Q30, Q39 using Fugene (4:1 ratio with DNA). Cells were treated with PBS or 0.25mM Sodium arsenite for 1 hour followed by 50*μ*g/mL emetine for 15 minutes.

### Puromycinylation Assay

Twenty hours post transfection primary cortical neurons expressing mScarlet-Ataxin-2 Q22, Q30, Q39, Q56, *Δ*Q or control RFP plasmid were treated with DMSO or 100*μ*g/mL cycloheximide for 30 minutes (51). O-propargyl-puromycin (OPP) was added at a concentration of 20 µM and incubated for 30 minutes. Neurons were rinsed once with prewarmed PBS and fixed in 4%PFA / 4% Sucrose in PBS. Click reaction was carried out with Click-iT Plus OPP Alexa Fluor 647 Protein Synthesis Assay kit (Thermo Fisher C10458) according to the manufacturer’s instructions, followed by immunofluorescence as described below.

### Immunofluorescence

HEK293 cells or mouse primary cortical neurons (DIV10-12) were fixed in PBS containing 4% PFA/4% sucrose for 12 minutes, washed three times in PBS, and quenched with 0.1 M glycine in PBS. Fixed neurons were permeabilized in 0.1% Triton X-100 in PBS for 12 minutes and blocked for 1 hour in PBS containing 5% normal goat serum, 1% BSA, and 0.1% Triton X-100. Samples were placed in a humidified chamber and incubated overnight at 4°C with primary antibodies, including rabbit anti-TDP-43 (Proteintech 1:300) or rat anti TDP-43 (R&D Systems, 1:300), rabbit anti-G3BP (Abcam, AB181150, 1:200), mouse anti-Ataxin-2 (BD Bosciences, 1:200), chicken anti-Tau (Synaptic Systems, 1:1000), mouse anti-FMRP (EMD Milipore, MAB2160, 1:500), and PABP (Abcam, AB21060, 1:1000). After five 10-minute washes in PBS, the samples were incubated in secondary antibody conjugated with Alexa Fluor dyes (Invitrogen, 1:200) for 1 hour at room temperature (RT), and then washed 5 times for 10 minutes each in PBS. Samples were then mounted in ProLong Gold (Life Technologies) and imaged either on an inverted NikonTi microscope with apochromat 60×1.49 NA oil-immersion objective with a PerkinElmer UltraVIEW VOX spinning disk confocal system and C9100-50 EMCCD camera (Hamamatsu) or on a Zeiss LSM 800 Axio Observer.Z1/7 Airyscan laser scanning confocal microscope with a Plan-Apochromat 63x/1.40 oil DICM27 objective.

### Single molecule FISH (smFISH)

smFISH protocol was adapted from (89). Briefly, rat cortical neurons (DIV8-10) were rinsed three times with PBS containing 5mM MgCl_2_ (5mM MgCl_2_/PBS) and fixed in PBS containing 4% PFA and 4% sucrose for 15 minutes. Neurons were treated with 0.1 M glycine solution made in 5mM MgCl_2_/PBS for 15 minutes, followed by three washes with 5mM MgCl_2_/PBS at RT for 5 minutes. Neurons were dehydrated in 50% ethanol, followed by 70% ethanol at RT for 2 minutes each. Neurons could be stored at this stage at 4°C for up to 1 week. Neurons were rehydrated sequentially, followed by equilibration in 5mM MgCl_2_/PBS at RT and incubation in 1X SSC [20X SSC (Invitrogen AM9763) diluted in Nuclease free water (AmericanBio, AB02128)] solution at RT for 10 minutes. For blocking, neurons were treated with 2X SSC solution containing 15% formamide (Ambion, AM9342, AM9344) and 5mg/ml ultrapure BSA (Invitrogen, AM2616) for 30 minutes at RT. Neurons were then incubated in pre-hybridization buffer [1X SSC solution containing 5% Dextran sulfate (Sigma, D8906), 10mg/ml ultrapure BSA, 15% formamide, 10mM ribonucleoside vanadyl complex (NEB, S1402S), 2*μ*g salmon sperm DNA (Invitrogen, 15632-011), 2*μ*g *E. coli* tRNA (Sigma-Aldrich, 10109541001) and 0.05X PBS] at 37°C for 90 minutes. After pre-hybridization, neurons were incubated with the pre-hybridization buffer containing *ACTB* probes labelled with Quasar 670 dye (Stellaris RNA FISH probes, Biosearch Technologies) at a final concentration of 500 nM for 20 hours at 37°C to label *β*-actin mRNA. To remove unbound probes, neurons were washed twice with 2X SSC buffer containing 15% formamide at 37°C for 30 minutes, followed by three brief washes with 2X SSC and three 5 minute washes with 2X SSC at RT on rocker. Samples were then mounted in ProLong Diamond antifade reagent (Invitrogen P36961) and allowed to cure overnight at RT. Images were captured using an inverted NikonTi microscope with apochromat 60×1.49 NA oil-immersion objective with Perkin Elmer UltraVIEW VOX spinning disk confocal system.

### SunTag Assay

SunTag assay was carried out as outlined previously (59, 90), with a few modifications. Rat cortical neurons (DIV9) were co-transfected with (1) mScarlet-TDP-43, mScarlet-Ataxin-2 of varying polyglutamine lengths, or control plasmid (RFP); (2) OsTIR1-IRES-scFV-sfGFP; and (3) pUbC-FLAG-24xSuntagV4-oxEBFP-AID-baUTR1-24xMS2V5-Wpre plasmids in a 1:1:1 ratio. Sixteen hours post-transfection, neurons were treated with indoleacetic acid (500*μ*g/ml, Sigma Aldrich I3750) for 5-6 hours to activate auxin induced degradation of mature Suntag-*β*-actin protein. To visualize mRNA, smFISH was performed using SunTag-V4 probes (59) labeled with Quasar 670 dye (Stellaris RNA FISH probes, Biosearch Technologies), as described above. As a control during validation of SunTag assay, neurons were treated with 100ug/mL of puromycin for 90 minutes at 37°C prior to fixation, as described (90)

### Live Cell Microscopy

Live imaging of cortical cultures expressing fluorescently-tagged TDP-43, Ataxin-2, or control cDNA constructs was performed in Hibernate E (Brainbits) supplemented with 2% B27 and 2mM GlutaMAX, in a temperature-controlled chamber mounted on an inverted NikonTi microscope with apochromat 60x 1.49 NA oil-immersion objectives; images were acquired on a PerkinElmer UltraVIEW VOX spinning disk confocal system equipped with an Ultraview Photokinesis (PerkinElmer) unit and a C9100-50 EMCCD camera (Hamamatsu) controlled by Volocity software (PerkinElmer). Axons and dendrites were identified based on morphologic criteria (91, 92), and images were captured at a rate of 0.5 frames per second for 5 minutes.

Fluorescence recovery after photobleaching (FRAP) experiments were performed using the Ultraview Photokinesis unit which was calibrated prior to each experiment to ensure tightly localized half-bleach, as described (33, 93). eGFP-TDP-43 was photobleached using the 488 nm laser at 50% power for 15 cycles, 1 ms spot period within a region of interest (0.25 × 0.25 *μ*m). Pre-bleach images were captured at 0.5 frames per second for 8 s, and subsequent to photobleaching, images were obtained at 1 frame per second for 120s.

### Image and Data Analysis

All image processing and analysis were performed using ImageJ/Fiji (94) and MATLAB R2019a or later (MathWorks). For Ataxin-2 and TDP-43 motility analysis, kymographs were prepared in Fiji and analyzed using custom MATLAB programs (33, 95) to calculate net and cumulative distance, net velocities, run lengths, fraction of “motile” and “non-motile” TDP-43 or Ataxin-2, and fraction of anterograde and retrograde transport. Condensates positive for TDP-43, without detectable Ataxin-2 were classified as TDP-43 positive (+); condensates positive for Ataxin-2 without detectable TDP-43 were classified as Ataxin-2 (+); and those condensates containing both TDP-43 and Ataxin-2 were classified as double-positive [TDP-43 (+) Ataxin-2 (+)]. Condensates that underwent long range directional transport, defined as net displacement of ≥10 *μ*m in 5 min or less, were classified as “motile”. Motile condensates were subdivided further by direction of net transport (anterograde or retrograde). Condensates that were transported <10 *μ*m were considered “non-motile”; this group was further broken down into stationary and oscillatory granules. Stationary TDP-43 granules were defined by < 5 *μ*m cumulative displacement, and oscillatory granules displayed > 5 *μ*m cumulative displacement. Proximal axon was defined as the portion of axon within 50 *μ*m of the soma, and mid axon was defined as the portion of axon >50 *μ*m from the cell body and >50 *μ*m from the axon terminal.

Time-lapse images of TDP-43 and Ataxin-2 axonal transport were denoised using a custom-written MATLAB pipeline that attenuated pixels using a binary mask, smoothing and attenuation thresholds. The pipeline set a noise-threshold, demarcating signal from noise, as a percentage of the maximum luminance from each frame. For example, a noise-threshold of 0.3 would create a binary mask where any pixel with a value less than 30% of the maximum luminance would be considered noise under the assumption that noise pixels captured less fluorescence than the targeted molecules that were emitting light and thus were separable based on luminance values. Once the binary mask was created, noise-pixels in that mask were attenuated by choosing a scaler value between 0 and 1 and multiplying each noise pixel by that value. A gaussian filter was then used on the mask to smooth the transition between noise and signal. Finally, using the chosen thresholds, the mask was created and applied to each frame and exported as an uncompressed AVI file. The code for this pipeline can be found on GitHub: https://github.com/dcalbick/DenoiseGUI.

For FRAP analysis, the mean fluorescence intensities from three regions of interest (ROIs) were obtained: 1) photobleached granule, 2) a non-bleached region (used to correct for overall photobleaching), and 3) background. A background subtraction was performed, the intensities of the photobleached ROI were normalized from 0 to 1 (pre-photobleach intensity = 1) and corrected for overall photobleaching. Corrected normalized fluorescence intensities were then plotted as a function of time, and curves were fit to a single exponential equation, I(t) = A(1-exp(-t/*τ*)), where I is normalized fluorescence intensity, t is time, *τ* is the characteristic time constant, and A is mobile fraction (96).

Quantification of TDP-43 and Ataxin-2 colocalization, or colocalization with polyA mRNA, puromycin, FMRP, G3BP1, or PABPC was performed using the stringent objects-based JaCoP plugin in Fiji (45). For each channel, a single slice was extracted from each z-stack, the axon was straightened, and the image was converted to 8 bit. Following thresholding, co-incidence of the centers of mass was used to identify double positive objects. For the analysis of *β-actin* mRNA fluorescence intensity along the mid axon and in the soma of each neuron, mean gray values were obtained within a defined region of interest (ROI) in the soma and in the mid axon. In the puromycinylation assay, the number of puromycin puncta per micron along the axon was quantified in Fiji by thresholding and using the analyze particles function. The number of puromycin puncta was divided by the length of the axonal segment analyzed to obtain the number of puromycin puncta/*μ*m.

SunTag and smFISH images were analyzed using FISH Quant (FQ) as described (54, 61). Briefly, Z-stack of confocal images of neuronal soma or dendrites were uploaded to FQ. After filtering images for background subtraction, Ataxin-2, TDP-43, scFV-sfGFP and mRNA signals were fit to a 3D Gaussian to determine the coordinates of puncta in each channel. The imaging parameters for these analyses were chosen to visualize only the granular signals. The intensity and the width of the 3D Gaussian were thresholded to exclude autofluorescent particles and non-specific signals: thresholds used for each channel were: >3000 for sfGFP; >1000 for Ataxin-2 or TDP-43; and >1500 for mRNA. For colocalization analysis and to quantify active translation sites where sfGFP SunTag puncta colocalized with mRNA, a custom MATLAB script was written which calculated the fraction of SunTag spots within 300 nm of mRNA, as described (54). The fraction of actively translating mRNA was defined as the number active translation sites divided by the total number of mRNA puncta. To quantify active translation sites associated with Ataxin-2 or TDP-43, triple colocalization events within 300nm from each other between all three channels were calculated.

### Statistical methods

Statistical tests were performed in GraphPad Prism, and curve fitting was performed in MATLAB R2019a or later. A Student’s *t* test (for normally distributed data) or Mann–Whitney *U* test (for non-normally distributed data) was used to compare two groups, and one-way analysis of variation with Tukey’s post hoc test was used to compare multiple groups of normally distributed data. The Kruskal– Wallis test with Dunn’s correction was performed to compare multiple groups of non-normally distributed data. The Kolmogorov–Smirnov test was used to compare frequency distributions. Box and whisker plots show the median (bar) and interquartile range (box), with whiskers extending from the fifth to 95th percentile. Sample sizes were chosen based on reported data from similar studies in the literature for the neuronal live-imaging field. Figure legends state *n*, the number of neurons or granules per condition; *N* represents the number of times the experiment was independently repeated.

