## Supplementary material for "Ataxin-2 polyglutamine expansions aberrantly sequester TDP-43, drive ribonucleoprotein condensate transport dysfunction and suppress local translation": SI Figures and Tables

**Supplementary Figure 1. Colocalization of endogenous TDP-43, Ataxin-2 and FMRP in a subset of axonal RNP condensates. (A-C)** Immunofluorescence for endogenous TDP-43 and Ataxin-2 in rodent primary cortical neurons (DIV 9-10). Representative images of different neurons are shown in A and B; the boxed area in panel B is enlarged in C. TDP-43 (green), Ataxin-2 (magenta), Tau (blue). Scale bars: **(A)** and **(B)** = 5 µm, **(C)** = 2.5 µm. **(D)** Colocalization of TDP-43 and Ataxin-2 with neuronal RNP transport granule marker, FMRP. Scale bar = 2.5 µm. **(E)** TDP-43 and Ataxin-2 colocalize with polyA mRNA to a similar extent; two-tailed t-test, (n=26 neurons for each condition, *N=3*) n.s., not significant.

**Supplementary Figure 2. Exogenous expression of fluorescently tagged wild type Ataxin-2 and/or TDP-43 shows similar cellular localization and expression pattern as endogenous Ataxin-2 and TDP-43 proteins. (A)** mScarlet-Ataxin-2 Q22 in the soma and **(B)** punctate expression along the axon of a primary cortical neuron co-expressing a control plasmid (LAMP1) mirrors endogenous expression seen in Figure 1 and Supplementary Figure 1. Scale bars: **(A)** soma = 3 µm, **(B)** axon = 2.5 µm. **(C)** eGFP-TDP-43 co-expressed with LAMP1 exhibits punctate expression in the axon, similar to endogenous TDP-43; scale bar = 2.5 µm. **(D)** Co-expression of eGFP-TDP-43 with mScarlet-Ataxin-2 Q22 shows punctate expression in the axon, similar to that seen with immunofluorescence for endogenous TDP-43 and Ataxin-2 in Figure 1; scale bar = 2.5 µm. **(E, F)** Kymographs show representative tracks for mScarlet-Ataxin-2 Q22 and eGFP-TDP-43 when each is co-expressed with LAMP1 control plasmid. Ataxin-2 condensates are mostly stationary; a similar motility pattern is seen when mScarlet-Ataxin-2 Q22 and eGFP-TDP-43 are expressed together (compare with Figure 2A). Scale bar = 30 sec, 2.5 µm.

**Supplementary Figure 3. RNP condensates positive for Ataxin-2 frequently associate with puromycin**. **(A)** Puromycinylation assay was used to visualize nascent peptide chains, followed by immunofluorescence for endogenous TDP-43, Ataxin-2, and tau in wild type primary cortical neurons. Neurons were pre-treated with cycloheximide or DMSO, followed by a 30 min pulse of O-propargyl-puromycin. Neurons were immunostained for endogenous TDP-43 (green), Ataxin-2 (magenta) and tau (blue). Scale bar = 3 µm. Click chemistry reaction was used to visualize puromycin incorporation into nascent polypeptide chains. **(B)** Ataxin-2 frequently colocalizes with puromycin in contrast to TDP-43,. ****$p<0.0001$, Mann-Whitney test, (n=15 neurons per condition, *N=3*). **(C)** Validation of the SunTag reporter assay with puromycin treatment. Puromycin abolishes bright sfGFP puncta (translation sites) and results in diffuse sfGFP signal (green) in the soma and dendrites of a primary cortical neuron; mRNA signal (magenta) is not affected. **(D)** A greater fraction of translating mRNA was observed in proximal dendrites of neurons expressing Ataxin-2 Q22 compared to TDP-43. ****$p<0.0001$, Mann-Whitney test. (Ataxin-2 n=26, TDP-43 n=24 neurons, *N=3*).

**Supplementary Figure 4. Ataxin-2 polyglutamine expansions sequester TDP-43 in stress granules and aggregates**
**(A, B)** HEK293T cells expressing control plasmid (pCMV-myc) or Ataxin-2 of varying polyglutamine lengths were treated with PBS or 25mM $NaAsO_2$ for 90 min, followed by immunofluorescence for G3BP (yellow), Ataxin-2 (magenta), TDP-43 (green) and DAPI stain (blue), TDP-43 zoomed-in panel represents magnified inset. Endogenous TDP-43 shows enhanced mislocalization to the cytoplasm in the presence of Ataxin-2 polyQ expansions in both control and oxidative stress conditions; scale bars = 10 µm. **(C)** Cytoplasmic TDP-43 aggregates $>20 \mu m^2$ form significantly more in cells expressing Ataxin-2 Q30 and Q39 compared to Q22 in both PBS and $NaAsO_2$ conditions, ****$p<0.0001$, Kruskal-Wallis test. **(D, E)** The integrated density of TDP-43 and G3BP within stress granules is higher in cells expressing Ataxin-2 Q30 or Q39 under control and oxidative stress conditions, Kruskal-Wallis test,

***p<0.0005, ****p<0.0001. **(F)** Following oxidative stress, Ataxin-2 Q22 incorporates into stress granule condensates that are predominantly TDP-43 (+) G3BP (+) double positive or G3BP (+). Ataxin-2 Q30 and Q39 incorporate into TDP-43 (+) G3BP (-) aggregates and stress granules. ****p<0.0001, 2-way ANOVA with Tukey's post-test (n=150 cells per condition, *N=3*).

**Supplementary Figure 5. Polyglutamine expansions (polyQ30, polyQ39) are not sufficient to sequester TDP-43 or G3BP1 in Ataxin-2 (+) condensates. (A)** Fluorescently-tagged polyQ30 and polyQ39 control constructs are unable to recruit TDP-43 into double positive condensates (Kruskal-Wallis test, n=20 neurons per condition, *N=3*), ns p>0.05). **(B)** Retrograde run lengths of motile and oscillatory TDP-43 condensates are not affected by the presence of Ataxin-2 Q22 or Q30. In neurons expressing Ataxin-2 Q39, TDP-43 (+) condensates show reduced retrograde run lengths, whether or not Ataxin-2 Q39 is in the condensate (*p<0.05; Kruskal-Wallis test, n=17-92 runs per condition from n=11-17 neurons, *N=3*). **(C)** The fraction of Ataxin-2 or **(D)** endogenous TDP-43 colocalizing with endogenous G3BP1 is significantly increased in neurons expressing Ataxin-2 Q30 or Ataxin-2 Q39, compared to neurons expressing Ataxin-2 Q22 or a control plasmid with fluorescently-tagged Q30 or Q39 tract (****p<0.0001, Kruskal-Wallis test; n=24, *N=3*).

**Supplementary Figure 6. Ataxin-2 polyglutamine expansions repress translation. (A, B)** Puromycinylation assay was performed with OPP, followed by click-chemistry reaction to visualize puromycin incorporation in primary cortical neurons expressing different Ataxin-2 polyQ variants or control plasmid. Reduced puromycin puncta are seen along the axon (no. of puromycin puncta/µm) in neurons expressing Ataxin-2 with polyQ tract of Q30 or longer, compared to controls; scale bar = 3 µm (***p<0.001, Kruskal-Wallis, n=15-16 neurons per condition, *N=3*). **(C, D)** SunTag reporter assay shows similar number of reporter mRNA in (**C**) the soma and (D) dendrites **D)** in neurons expressing control RFP plasmid or Ataxin-2 polyQ variants, indicating similar expression levels between conditions (ns p>0.05, Kruskal-Wallis test; n=24, *N=3*).

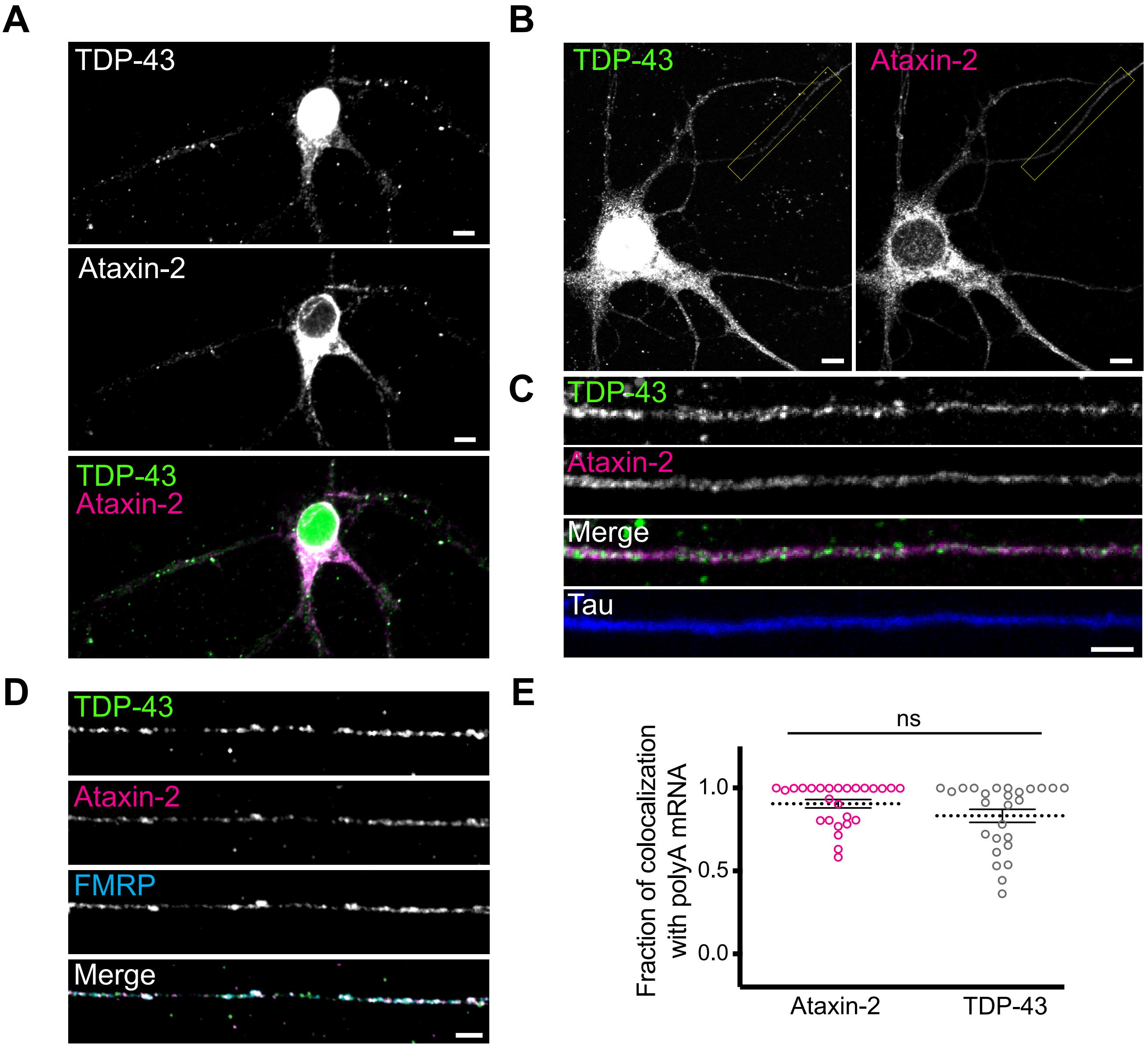


for supplementary figure 1
A, B: scale bar = 5 um
C: scale bar = 2.5 um
D: scale bar = 25.5 pix

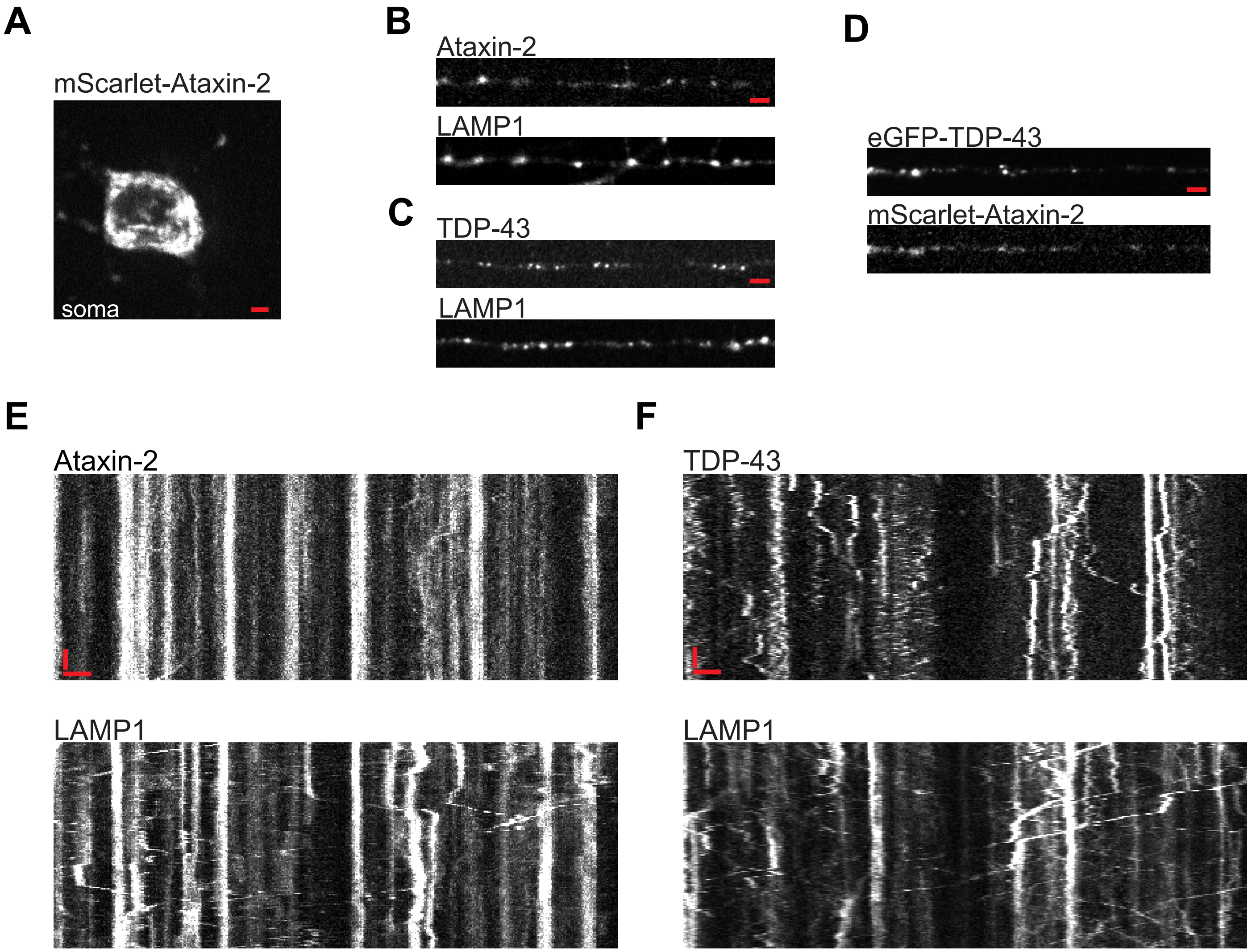
A
mScarlet-Ataxin-2
soma
B
Ataxin-2
LAMP1
C
TDP-43
LAMP1
D
eGFP-TDP-43
mScarlet-Ataxin-2
E
Ataxin-2
LAMP1
F
TDP-43
LAMP1

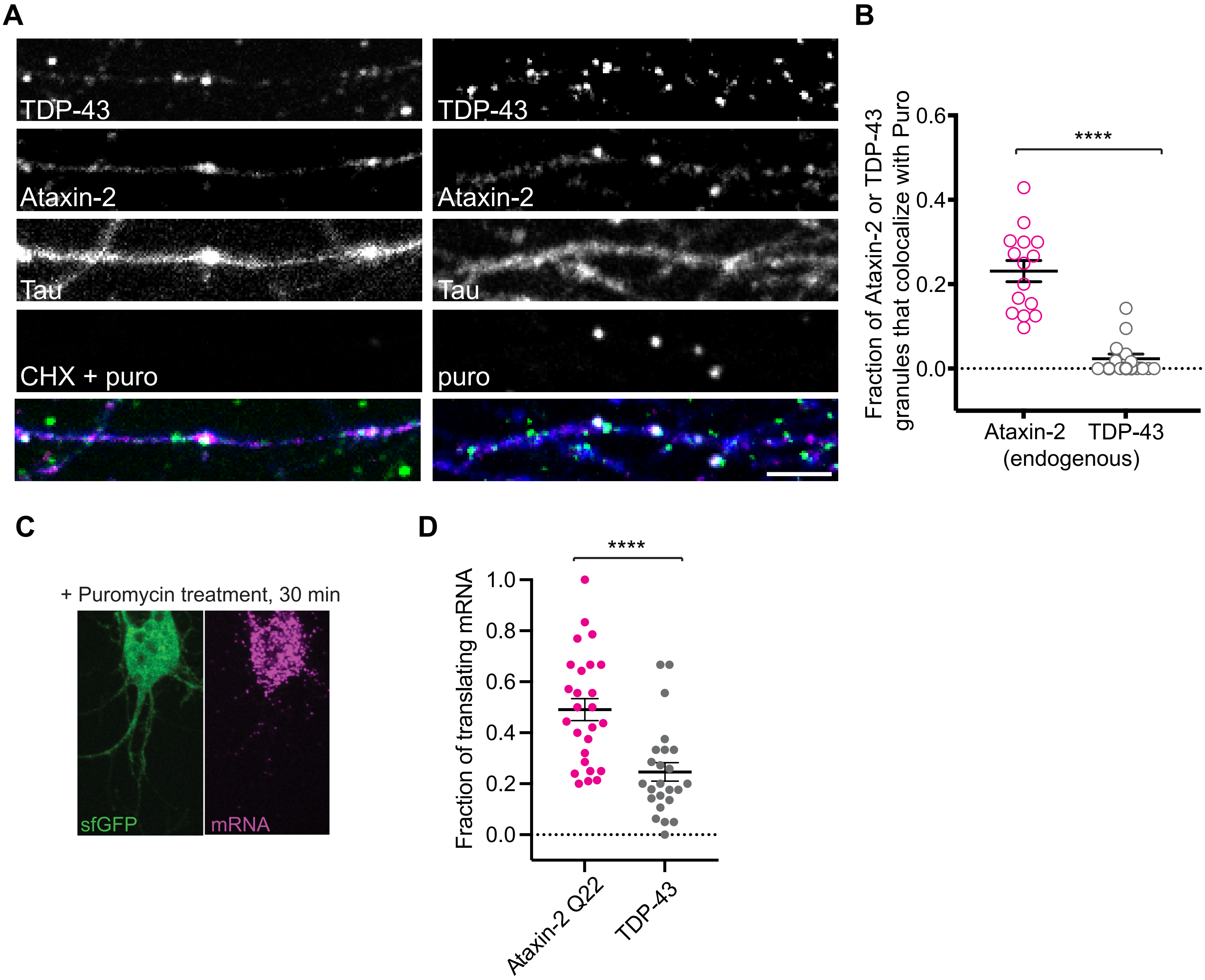
A
TDP-43
Ataxin-2
Tau
CHX + puro
TDP-43
Ataxin-2
Tau
puro
B
Fraction of Ataxin-2 or TDP-43 granules that colocalize with Puro
0.6
0.4
0.2
0.0
****
Ataxin-2
TDP-43
(endogenous)
C
+ Puromycin treatment, 30 min
sfGFP
mRNA
D
Fraction of translating mRNA
1.0
0.8
0.6
0.4
0.2
0.0
****
Ataxin-2 Q22
TDP-43

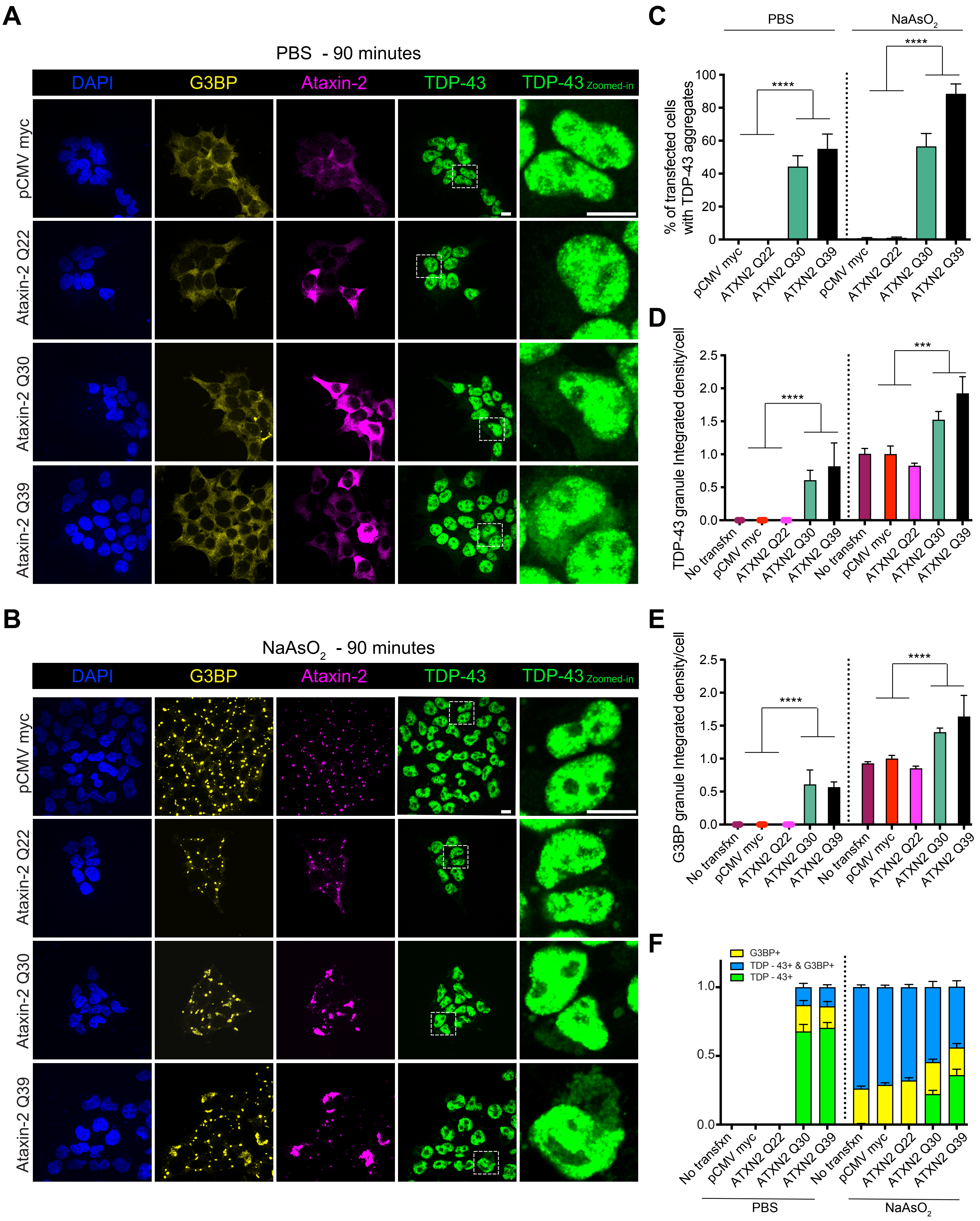
A
PBS - 90 minutes
DAPI
G3BP
Ataxin-2
TDP-43
TDP-43 Zoomed-in
pCMV myc
Ataxin-2 Q22
Ataxin-2 Q30
Ataxin-2 Q39
B
NaAsO2 - 90 minutes
C
PBS
NaAsO2
% of transfected cells with TDP-43 aggregates
pCMV myc
ATXN2 Q22
ATXN2 Q30
ATXN2 Q39
D
TDP-43 granule Integrated density/cell
No transfxn
E
G3BP granule Integrated density/cell
F
G3BP+
TDP - 43+ & G3BP+
TDP - 43+

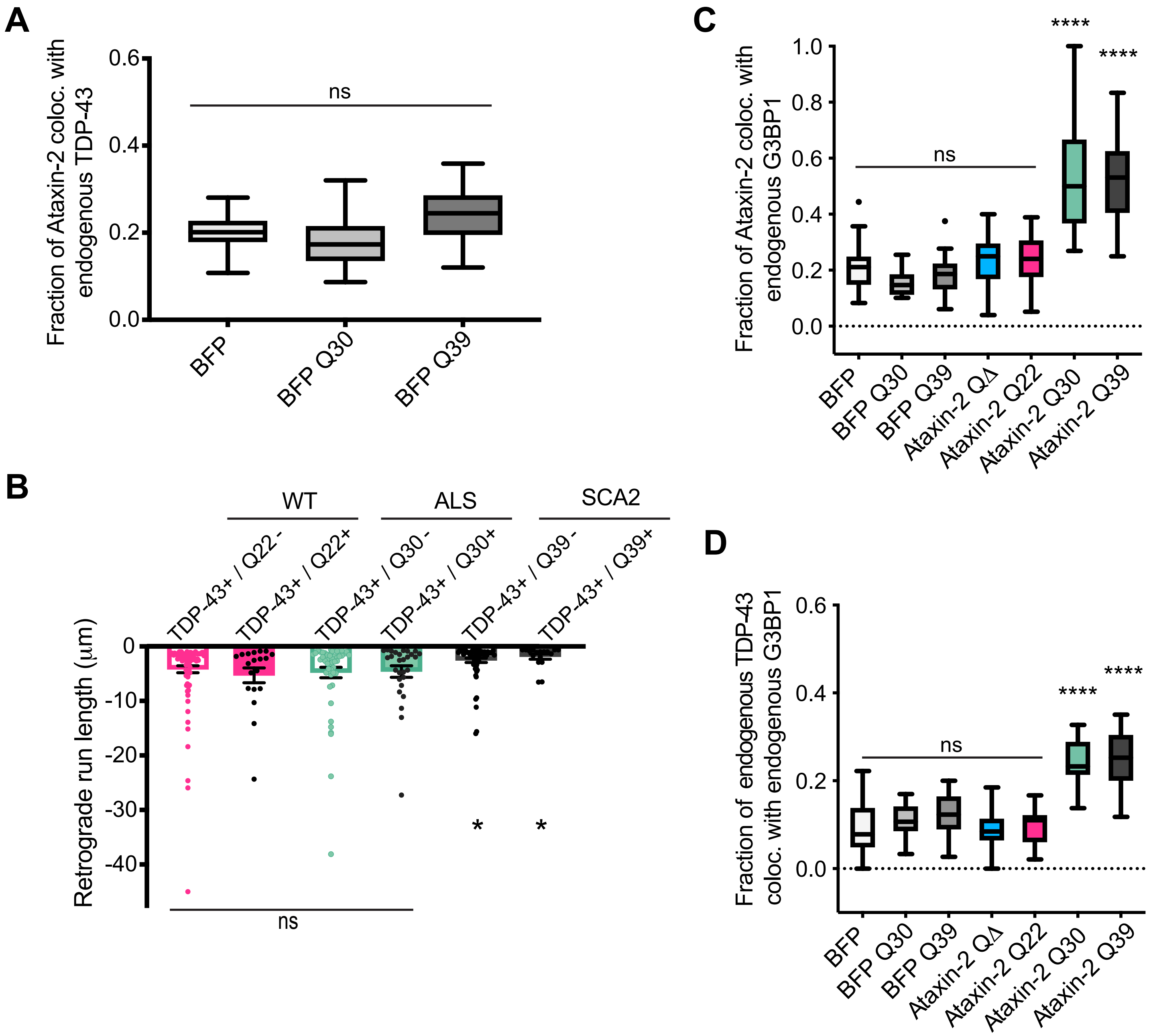

A
Fraction of Ataxin-2 coloc. with endogenous TDP-43
0.6
0.4
0.2
0.0
ns
BFP
BFP Q30
BFP Q39
B
WT
ALS
SCA2
TDP-43+ / Q22 -
TDP-43+ / Q22+
TDP-43+ / Q30 -
TDP-43+ / Q30+
TDP-43+ / Q39 -
TDP-43+ / Q39+
Retrograde run length (μm)
0
-10
-20
-30
-40
*
*
ns
C
Fraction of Ataxin-2 coloc. with endogenous G3BP1
1.0
0.8
0.6
0.4
0.2
0.0
ns
****
****
BFP
BFP Q30
BFP Q39
Ataxin-2 QΔ
Ataxin-2 Q22
Ataxin-2 Q30
Ataxin-2 Q39
D
Fraction of endogenous TDP-43 coloc. with endogenous G3BP1
0.6
0.4
0.2
0.0
ns
****
****
BFP
BFP Q30
BFP Q39
Ataxin-2 QΔ
Ataxin-2 Q22
Ataxin-2 Q30
Ataxin-2 Q39

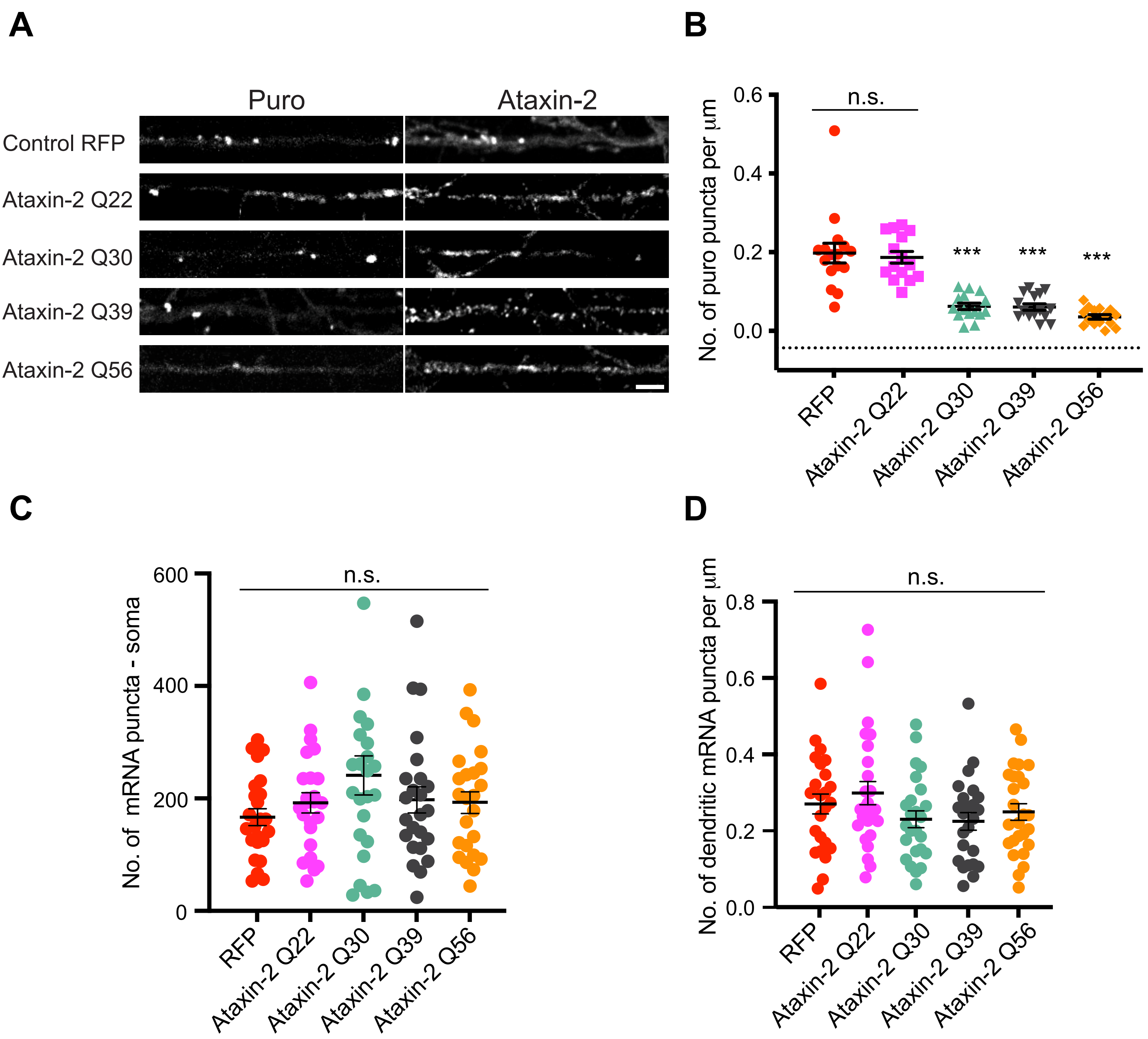

A
Puro
Ataxin-2
Control RFP
Ataxin-2 Q22
Ataxin-2 Q30
Ataxin-2 Q39
Ataxin-2 Q56
B
No. of puro puncta per µm
n.s.
***
RFP
Ataxin-2 Q22
Ataxin-2 Q30
Ataxin-2 Q39
Ataxin-2 Q56
C
No. of mRNA puncta - soma
n.s.
D
No. of dendritic mRNA puncta per µm
n.s.

**Table. Axonal TDP-43 or Ataxin-2 condensate FRAP recovery time constants, related to Figure 2 and Figure 6.**

| | **τ (s) with 95% CI** | **MATLAB Curve fitting** |
|---|---|---|
| **Ataxin-2** | 25.5<br>(21.3, 31.8) | Single exponential<br>$I(t) = A(1-exp(-t/\tau))$<br>A = 0.28 (.26, .30) |
| **TDP-43 (negative for Ataxin-2 Q22** | 21.1<br>(19.9, 22.5) | Single exponential<br>$I(t) = A(1-exp(-t/\tau))$<br>A = 0.58 (.56, .59) |
| **TDP-43 RNP condensates with Ataxin-2 Q22**<br>**half bleach** | 13.6<br>(12.1, 15.7) | Single exponential<br>$I(t) = A(1-exp(-t/\tau))$<br>A = 0.51 (.48, .55) |
| **TDP-43 RNP condensates with Ataxin-2 Q30**<br>**half bleach** | 17.7<br>(16.0, 19.9) | Single exponential<br>$I(t) = A(1-exp(-t/\tau))$<br>A = 0.36 ( .34 .38) |
| **TDP-43 RNP condensates with Ataxin-2 Q39**<br>**half bleach** | 33.1<br>(29.0, 38.5) | Single exponential<br>$I(t) = A(1-exp(-t/\tau))$<br>A = 0.30 (.28, .31) |

I = Intensity, t = time

Table 2. Primers

| Primer Name | Sequence |
| --- | --- |
| **ΔQ Fwd** | ccgccgcctgcagctgccaatgtccgca |
| **ΔQ Rev** | agctgcaggcggcgggggcttcagcgacatggt |
| **Q56 Fwd** | caacaacaacaacaacaacaacaacaacaacaacaaccgccgccgcggctgc |
| **Q56 Rev** | cggcggcttcttcttcttcttcttcttcttcttcttcttctgctgctgctgctg |
| **Q30 and Q39 Fwd** (for BFP Q30/ Q39) | aaagatctaccatgtcgctgaagccc |
| **Q30 and Q39 Rev** (for BFP Q30/ Q39) | aaggtaccgggcttgcggacattggc |
